# Single-cell molecular profiling using ex vivo functional readouts fuels precision oncology in glioblastoma

**DOI:** 10.1101/2022.10.17.512525

**Authors:** Dena Panovska, Pouya Nazari, Basiel Cole, Pieter-Jan Creemers, Marleen Derweduwe, Lien Solie, Sofie Van Gassen, Annelies Claeys, Tatjana Verbeke, Yvan Saeys, David Van Der Plancken, Francesca M. Bosisio, Eric Put, Sven Bamps, Paul M. Clement, Michiel Verfaillie, Raf Sciot, Keith L. Ligon, Steven De Vleeschouwer, Asier Antoranz, Frederik De Smet

## Abstract

**Background:** Functional profiling of freshly isolated glioblastoma cells is being evaluated as a next-generation method for precision oncology. While promising, its success largely depends on the method to evaluate treatment activity which requires sufficient resolution and specificity.

**Methods:** Here, we describe the ‘<u>pr</u>ecision <u>o</u>ncology by <u>s</u>ingle-cell <u>p</u>rofiling using *<u>e</u>x vivo* readouts of functi<u>o</u>nality’ (PROSPERO) assay to evaluate the intrinsic susceptibility of high-grade brain tumor cells to respond to therapy. Different from other assays, PROSPERO extends beyond life/death screening by rapidly evaluating acute molecular drug responses at single-cell resolution.

**Results:** The PROSPERO assay was developed by correlating short-term single-cell molecular signatures using CyTOF to long-term cytotoxicity readouts in representative patient-derived glioblastoma cell cultures (n=14) that were exposed to radiotherapy and the small-molecule p53/MDM2 inhibitor AMG232. The predictive model was subsequently projected to evaluate drug activity in freshly resected GBM samples from patients (n=34). *Here,* PROSPERO revealed an overall limited capacity of tumor cells to respond to therapy, as reflected by the inability to induce key molecular markers upon *ex vivo* treatment exposure, while retaining proliferative capacity, insights that were validated in PDX models. This approach also allowed the investigation of cellular plasticity, which in PDCLs highlighted therapy-induced proneural-to-mesenchymal transitions, while in patients’ samples this was more heterogeneous.

**Conclusion:** PROSPERO provides a precise way to evaluate therapy efficacy by measuring molecular drug responses using specific biomarker changes in freshly resected brain tumor samples, in addition to providing key functional insights in cellular behavior, which may ultimately complement standard, clinical biomarker evaluations.

## Introduction

Glioblastoma (GBM), a uniformly lethal brain tumor, could greatly benefit from the availability of better diagnostics to match potential therapeutic options to the right patient subpopulation [1, 2]. Already for more than 15 years, the uniformed standard-of-care protocol for GBM has not been changed, despite extensive insights in inter-patient and intra-tumoral heterogeneity [1,3,4]. Major efforts over the past two decades have aimed at identifying better therapies for GBM, which so far did not lead to substantial changes in overall survival [5]. However, in spite of the overall inability to treat GBM, clinical trials often describe anecdotical or small groups of patients that did show a clinical response [6, 7], suggesting that methods to better identify these exceptional patients could lead to a potentially improved clinical management.

A functional precision oncology assay would be highly suited to fill this gap. However, in the particular case of GBM, such an assay should not only be able to map each tumor in great detail—given GBM’s high degree of heterogeneity [8]—it should also be applicable to every patient in either the newly diagnosed or recurrent setting, lead to actionable results within a relevant timeframe, and provide specific insights in drug activity. A variety of functional diagnostic (FD) tests are gaining traction to identify patient-tailored therapeutic options for GBM [9–12]. In these ongoing efforts, a primary focus is put on establishing patient-derived cultures [13] or short-cultured explants [14] which are subsequently screened in semi-high-throughput assays to determine chemo-sensitivity profiles (typically using life/death assays) with the goal to eventually correlate the obtained profiles to patients’ prognosis [9,10,15,16]. While of major interest, these assays (i) lack the ability to systematically generate culture models for most GBM patients (take rate of ∼30-60%) [17]; (ii) require several months to generate sensitivity profiles, while (iii) of insufficient scalability to test diverse treatment modalities.

In this proof-of-concept study, we developed the PROSPERO assay, a functional precision oncology (FPO) workflow for GBM that enables us to map drug activity at single-resolution in freshly collected tumor samples without the need for prolonged culturing/expansion. As opposed to other FPO assays, this was done by measuring fast, therapy-related biomarker changes using high dimensional, single cell profiling using mass-cytometry (CyTOF), as such going beyond life/death screening. This approach allowed us to also evaluate therapy-induced cellular plasticity, a feature that is of major importance in establishing treatment resistance in GBM. Importantly, the assay generated insights in drug activity within days following surgery and was compatible with samples obtained from both newly-diagnosed and recurrent patients. Finally, the identified functional drug-dependent signatures were also correlated to *in vitro* long-term cytotoxicity profiles in patient-derived models, as such generating a computational model to predict drug sensitivity in freshly collected patient samples. While full clinical evaluation is still needed, this study offers a cost/time effective functional precision medicine assay that could help to guide future clinical decision-making in the treatment of glioblastoma.

## Material and Methods

### GBM sample collection and ethical approval

Freshly resected high-grade glioma tumor samples were collected from the University Hospitals Leuven (Leuven, Belgium; IRB protocol S59804), Europaziekenhuizen (Brussels, Belgium; IRB protocol EC approval 05-02-2018), Jessa Ziekenhuis (Hasselt, Belgium, IRB B243201941451), and AZNikolaas (Sint Niklaas, Belgium; IRB protocol EC18021). Consent was given by all patients. Histopathological analysis and IDH mutation assessment were performed by a certified neuropathologist in all cases.

Patient-derived cell lines (PDCLs) were generated from tumor samples and characterized as previously described [18]. Cell lines generated in Leuven (between 2017-2019) ((according to IRB protocol S59804, LBT-numbers (Leuven, Belgium). Cell lines obtained from Dana-Farber Cancer Institute (BT-numbers (Boston, MA, USA)) in 2017 (IRB 10-417). Cells were authenticated by testing short tandem repeats using the PowerPlex Fusion System (DC2408, Promega). The most recent authentication was performed in July, 2019. An overview of the 14 PDCLs can be found in Supplementary Table 1.

### Drug treatments and cytotoxicity assays using AMG232 and RT

Short-term drug exposure and CyTOF sample preparation was performed on PDCLs and biopsy samples. Viability assays were performed only on PDCLs (see more details in Supplementary Methods).

### Mouse PDX experiments

All protocols involving work with live animals were reviewed and approved by the KULeuven Animal Ethics Committee under the protocol P211/2018, and experiments were conducted in accordance with the KULeuven animal facility regulations and policies. Orthotopic models were generated as previously described [19] and further explained in Supplementary Methods.

### CyTOF experiments

All steps were performed according to manufacturer’s instructions, unless indicated and outlined in Supplementary Methods.

### Data analysis

Tumor subtype annotation and identification of phenotypic patterns in PDCLs and patients’ samples. Different tumor cell types were identified following a consensus clustering strategy [20, 21] (see Supplementary Methods for more details).

### Identification of cell cycle phase

Cells were assigned to one of the four pre-defined cell cycle phases (G0G1, S, G2, M) after manual gating, as previously described [22].

### Unbiased probability modeling of drug responses in PDCLs using AUC and PLI

The induction of functional markers was evaluated for the different treatments and samples (PDCLs, tumor biopsies, PDX samples). Unbiased response modeling and correlation to AUC and PLI as endpoints are further explained in Supplementary Methods.

### Projection of the predictive model on patients’ cohort

These fine-tuned models were used to project the biopsy samples and give an estimation on their drug-response. The same approach was used using only the control samples and cell cycle.

### Mouse PDX experiments

All protocols involving work with live animals were reviewed and approved by the KULeuven Animal Ethics Committee under the protocol P211/2018, and experiments were conducted in accordance with the KULeuven animal facility regulations and policies. The data analysis is explained in Supplementary Methods.

### Code availability

All codes are provided on the github page, which will be made available upon publication of the manuscript.

## Results

### Study design

In this proof-of-concept study we tested the technical ability of the ‘<u>pr</u>ecision <u>o</u>ncology by <u>s</u>ingle-cell <u>p</u>rofiling using *<u>e</u>x vivo* readouts of functi<u>o</u>nality’ (PROSPERO) assay to capture the intrinsic capabilities of *ex vivo* treated glioblastoma cells to respond to therapeutic insults. To set up this workflow, various preclinical models were used, including patient-derived high-grade glioma stem cell lines (PDCLs) and mouse PDX models (Figure 1), following which the developed analysis pipeline was applied to freshly resected tumor samples from GBM patients treated at various Belgian hospitals. We therefore first aimed at building and optimizing an analysis pipeline for which we correlated long-term cytotoxicity profiles of PDCLs upon exposure to AMG232 or radiation therapy (RT), to early stage single-cell, phenotypic and functional signatures upon drug perturbation (Figure 1, Figure 2A). Based on the observed biomarker changes, a predictive model was generated, which was subsequently applied to determine drug activity in freshy resected GBM samples. By measuring single-cell drug perturbation signatures within hours following surgery, we could estimate the percentage of intrinsically sensitive/resistant tumor cells in each individual sample while simultaneously measuring therapy-induced proliferative and phenotypic transitions (Figure 1). As such, this strategy raises the specificity and spectrum of generated insights that go well beyond mere life/death screening.

**Fig 1.**
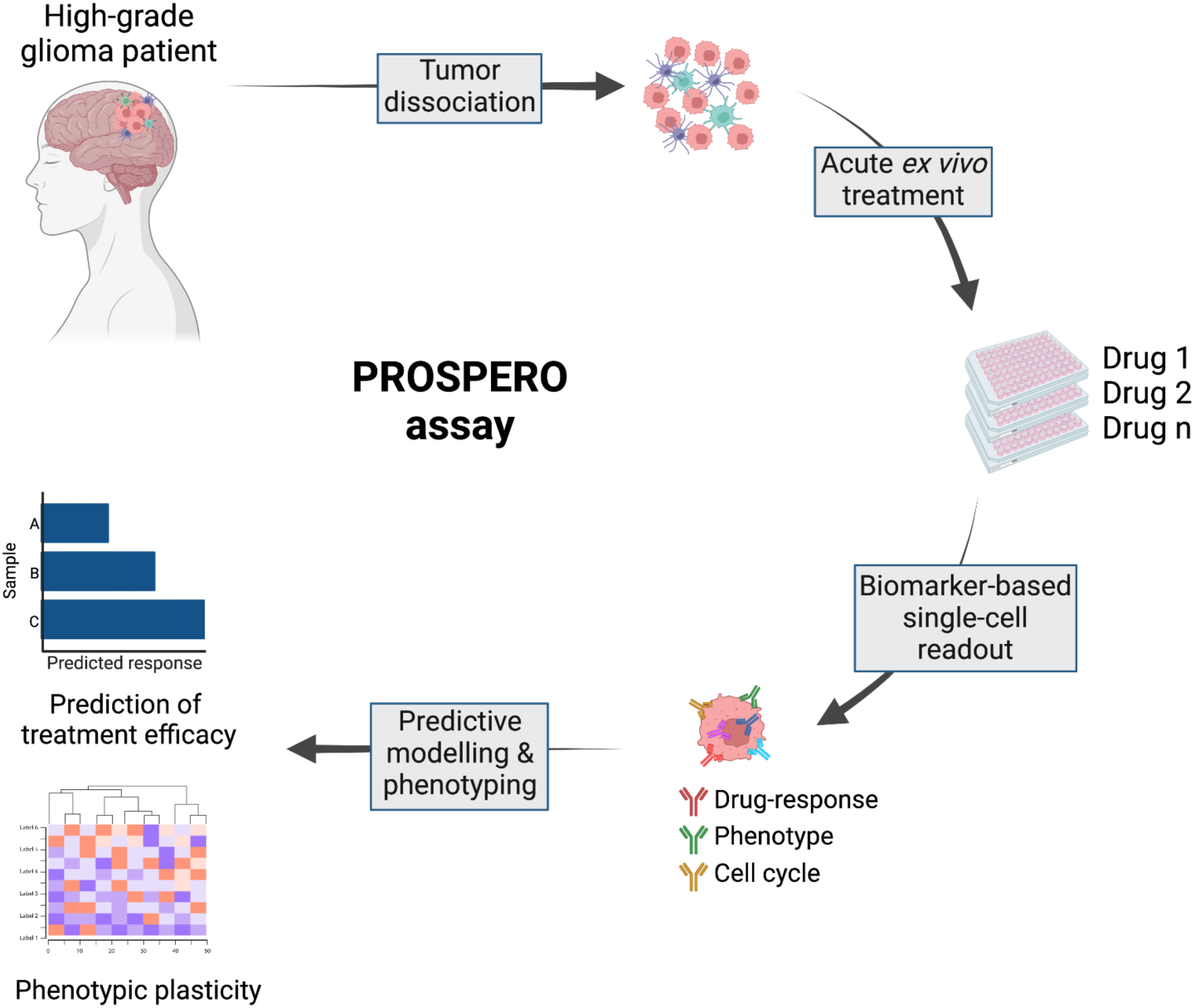
Setup of PROSPERO assay focusing on functional analysis

**Fig 2.**
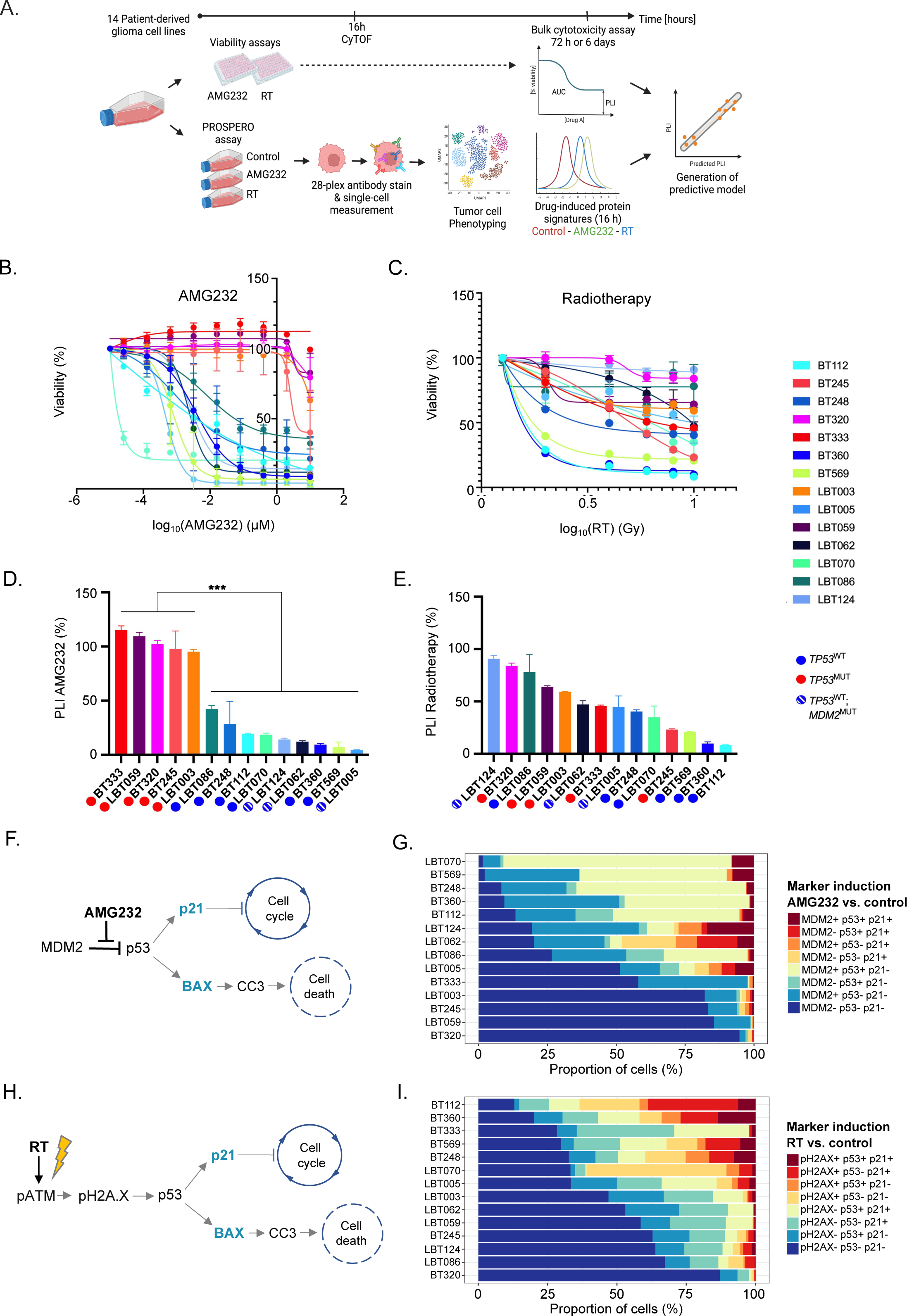
Mapping of bulk cytotoxicity profiles and single-cell drug heterogeneity in PDCLs. (a) PROSPERO workflow in PDCLs. (b, c) Dose-response curves representing the cell viability after AMG232 and RT, respectively. (d, e) Bar charts of PLIs after AMG232 (0-10μM for 72 hours) and RT (0-10 Gy for 6 days), respectively. The values represent mean ± SD of three (AMG232) or four (RT) replicates and were normalized to 100% assigned to the vehicle control for each assay. Overall responses of the models stratified by *TP53* status and assigned colored circles (*TP53*^MUT^-red; *TP53*^WT^-blue and *MDM2*^AMP^ – white/blue stripes). Wilcoxon rank sum test was used to calculate statistical significance between the *TP53*^WT^ and *TP53*^MUT^ groups (ns = not significant (P > 0.05); *P ≤ 0.05; **P ≤ 0.01; ***P ≤ 0.001; ****P ≤ 0.0001). (f) AMG232 targeting MDM2/p53 complex and downstream cellular effects. (g) Stacked barplots representation of drug-induced signatures across the PDCLs (n=14) ordered by response to AMG232 (from most to least responsive). (h) Molecular effects induced by RT. (i) Stacked barplots representation of drug-induced signatures across the PDCLs (n=14) ordered by response to RT (from most to least responsive)

### CyTOF enables simultaneous single-cell assessment of phenotype and treatment-induced molecular markers in PDCLs

The proposed concept of functional diagnostics for GBM consists of *ex vivo* treatment of freshly resected patients’ samples and using alterations of biomarkers’ expression as response readouts. Initially, to develop and standardize the required procedures and tools for the functional diagnostic platform, we selected a group of patient-derived *IDH*^WT^ GBM cell lines (PDCLs, n=14), exhibiting a typical range of genetic alterations commonly found across GBM patients (including deletion of *CDKN2A*; amplification of *CDK4/6, EGFR, MDM2/4*; mutations in *TP53, PTEN, PIK3CA, EGFR, MET* and *NF1*) [4], express primary pathological GBM markers (e.g. Sox2, GFAP, Olig2, and/or Nestin) and cover previously described GBM subtypes [1, 3], based on bulk expression measurements (Supplementary Figure 1A & B; Supplementary Table 1). The protein expression of several phenotypic markers was assessed by CyTOF analysis (Supplementary Figure 1C & D) and overall correlated well to bulk RNA sequencing (Supplementary Figure 1E). The discrepancy between RNA and protein levels of p53 in the PDCLs could be explained by the autoregulation of p53 levels by negative feedback loops and post-translational modifications of mutant p53, which enable p53 to escape MDM2-mediated degradation and accumulation in the nucleus [23–25].

Not only did these models closely capture GBM’s pathophysiology, they also allowed us to develop and optimize all procedures and tools necessary for a functional diagnostic platform, such as drug concentrations (for the selected therapies), length of treatment exposure and antibody panels. Then, we sought to perform long-term viability treatments in parallel to short-term acute treatments on the same models to ultimately correlate (early) protein expression changes (control vs treated) to long-term cell survival. Based on these insights, we trained a predictive algorithm of response (Figure 2A).

As a benchmark, we applied AMG232 [26], a selective MDM2 inhibitor, currently in clinical trial for GBM (ClinicalTrials.gov Identifier: NCT03107780). MDM2 is a negative regulator of the tumor-suppressor gene *TP53* [19] and AMG232 has been proven to robustly induce apoptosis and cell cycle arrest, specifically in *TP53*^WT^ cells [19]. We also exposed the PDCL pool to ionizing radiation treatment (RT), as part of the standard-of-care therapy for GBM patients [27]. As anticipated, AMG232 induced significantly higher cell death in *TP53*^WT^ compared to *TP53*^MUT^ cell lines (p=<0.0001; Figure 2B). Upon RT, we observed a wide range of sensitivities (from resistant to highly sensitive) for which the *TP53* mutational status alone nor any other measured genetic alteration seemed able to directly correlate to sensitivity (Figure 2C), which is in line with its less specific mode-of-action (MoA) [26].

These bulk dose-response analyses enabled the extrapolation of IC_50_, area under the curve (AUC) and plateau level of inhibition (PLI; a measure that reflects the percentage of non-responsive cells in a PDCL upon prolonged exposure) (Figure 2B-E; Supplementary Figure 2A-B; Supplementary Table 1). In both therapies, these readouts varied greatly across the *TP53*^WT^ pool, which suggests the presence of heterogeneous cell populations with diverse drug response capacity. Thus, we hypothesized that within each *TP53*^WT^ cell line there are populations which are either intrinsically resistant or at least unable to induce a full drug response. For the purpose of the downstream analyses and generation of the predictive model, we focused on PLI values as indicators of long-term drug tolerability at bulk level (Figure 2D & E).

Developing a reliable functional diagnostic assay to quantify the ability of GBM cells to elicit a genuine molecular drug response in phenotypically heterogeneous cell populations, implied the a-priori selection of markers of interest (Figure 2F & H, Supplementary Table 2). As such, we optimized a panel of antibodies based on previous knowledge regarding the molecular MoAs [26, 28] of both included therapies, as well as phenotype-specific features (ie GBM subtypes), as described by single-cell RNA-directed studies [1] (Supplementary Table 2 and 3). For AMG232, the molecular mechanism has been well characterized [29]: upon inhibition of the MDM2-p53 interaction, the p53 protein accumulates in the nucleus, and, if still functionally active, induces the expression of downstream targets including the cell cycle inhibitor p21/CDKN2A and/or the apoptosis-inducer BAX. From these, p21 causes cell cycle arrest through the inhibition of the cyclin dependent kinase CDK2, while activated BAX will lead to apoptosis commonly mediated through cleaved caspase-3 (CC-3) [30] (Fig. 2F). Ionizing radiation, on the other hand, typically causes DNA double strand breaks [31], which result in the early activation of ATM by phosphorylation (pATM) which in its turn (i) phosphorylates H2AX to assist in the repair the DNA damage and (ii) activates p53, resulting in the induction of the same downstream molecular pathway as described for AMG232 [30] (Fig. 2H). Based on these insights, we assembled a drug-related antibody panel targeting pATM, pH2AX, MDM2, p53, p21, BAX and CC-3.

To validate PRSOPERO’s capability in capturing drug response heterogeneity (intrinsically resistant subpopulations) we mixed, in various proportions, and treated two PDCLs (BT360 and BT333; Supplementary Materials) which harbor differential responsiveness. First, bulk cytotoxicity measurements of these mixtures demonstrated that the range of responsiveness profiles extending between the individual (“pure”) cultures followed the proportion of BT360 and BT333 in each mixture (Supplementary Figure 2C & D). The PROSPERO assay showed that the single-cell protein expression changes of the drug-related markers (Supplementary Figure 2E & F) were corresponding to the bulk viability profiles (Supplementary Figure 2C & D), highlighting the ability of PROSPERO to capture the proportion of unresponsive cells at single-cell resolution.

The induction of each of the treatment-dependent biomarkers was determined in every single cell using a statistical probability model, comparing biomarker expression upon treatment to a corresponding control sample that was included in every experiment and for every individual sample. This approach enabled us to rank the models from most to least responsive to the given therapies based on the ability of individual cells to induce (combinations of) biomarkers upon exposure to therapy (Figure 2G & I). Similarly, as in the dose-response assays, the models could be separated by the *TP53-*status: *TP53*^WT^ cells present larger numbers of responsive cells (upon AMG232) treatment, while this separation based on *TP53* status was not obvious upon RT.

### Unbiased modeling of drug responses in PDCLs identifies marker combinations which correlate to PLI

Next, we set out to generate an unbiased model to predict long-term cytotoxicity based on the acute induction of treatment-related biomarkers (Supplementary Figure 3). To this end, we correlated acute treatment-dependent signatures (measured at 16 hours) to dose-response cytotoxicity assays performed in parallel for each PDCL. Pseudotime analysis was performed to rank each PDCL along a continuum ranging from ‘no induction’ (low pseudotime value) to ‘full induction’ (high pseudotime value) for each possible marker combination, reflecting the response spectrum of the PDCL pool. Next, these pseudotime values were correlated (Pearson correlation/R) to the observed PLI/AUC, representing an unbiased approach to find the optimal marker combination with the best predictive power (Figure 3A & D, Fig. Supplementary Figure 4). Interestingly, the literature-guided marker combination for AMG232 treatment (MDM2/p53/p21; (R=-0.759)) and RT (pH2AX/p53/p21; (R=-0.380)) did not yield the best correlation to PLI, underscoring the importance of the unbiased approach (Supplementary Figure 4). Based on the correlation values and the law of diminishing marginal returns, the optimal (unbiased) marker to predict PLI after AMG232 treatment only required a functional measurement of p21 induction (R=-0.827, Figure 3A, Supplementary Figure 4B), while the optimal combination for RT treatment was determined by the induction of p21/pATM/pH2AX (R=-0.801, Figure 3D & E, Supplementary Figure 4D).

**Fig 3.**
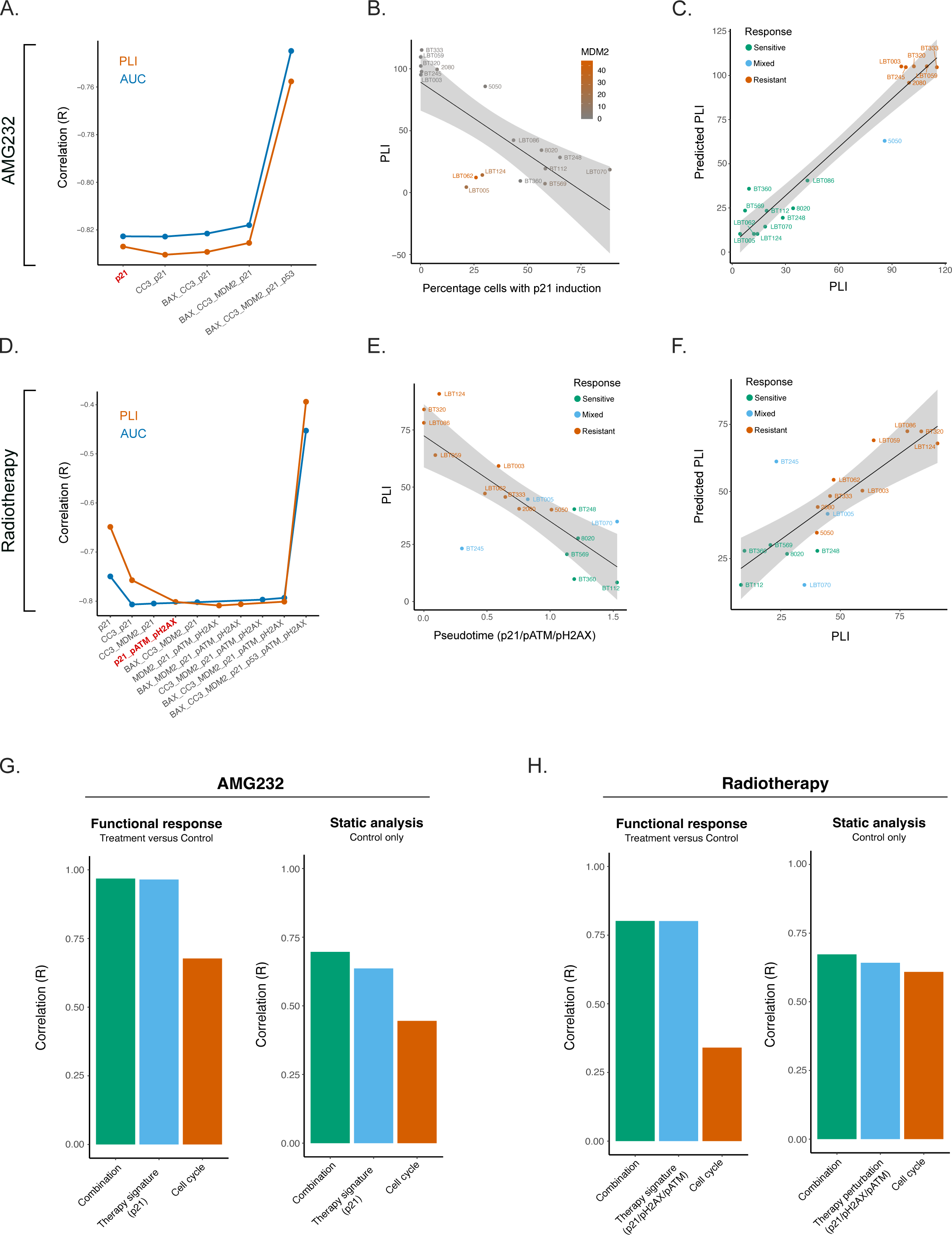
Unbiased probability modeling of AMG232 and RT responses. (a-c) AMG232 treated samples. (a) Pareto front of signature optimization correlated to AUC/PLI (R). The preferred signature with highest correlation (R) to PLI is highlighted in red (p21). (b) Pseudotime of the correlation between percentage of cells with p21 induction in each sample and PLI, highlighting *MDM2*^AMP^ cell lines. (c) Correlation between PLI and predicted PLI. (d-f) RT treated samples. (d) Pareto front of RT panel optimization correlated to AUC/PLI (R). The preferred signature with highest correlation (R) to PLI is highlighted in red (p21/pATM/pH2AX). (e) Correlation of the pseudotime to PLI. (f) Correlation between PLI and Predicted PLI. (g, h) Testing the prognostic capacity of each signature set (functional, cell cycle or combination of both) in drug-(left panels) and vehicle-treated cells (right panels)

Interestingly, the linearity of the correlation between the p21 induction value and the PLI for AMG232 treatment was skewed by *MDM2*^AMP^ PDCLs (LBT005, LBT062, LBT124), suggesting that a different probability model applies to these samples (Figure 3B). When *TP53*^WT^/*MDM2*^AMP^ cell lines are separated from the rest of the PDCL pool, an improved correlation could be achieved for PLI (Figure 3C; Supplementary Figure 5A & B; Supplementary Figure 6C & D) and AUC (Supplementary Figure 6A & B). Finally, we tested the correlation between the predicted PLI, resulting from the AMG232/RT response modeling and the actual PLI (Supplementary Figure 5C-F), which were following a linear distribution (Figure 3C & F).

### Therapy-induced cell cycle arrest of highly proliferative glioma cells is *TP53*-dependent

P21 represents an indicator of therapy responsiveness and a reliable biomarker for molecular readout upon AMG232/RT treatment [19, 28]. Therefore, we wanted to explore the downstream effects on the cell cycle caused by p21 upregulation.

For each PDCL, we firstly distinguished cells in different cell cycle phases and subsequently mapped the percentages of cells belonging to each treatment condition (Supplementary Figure 7; Supplementary Figure 8A & B). In general, both therapies were able to affect cell cycle progression to some extent. In the *TP53*^WT^ group, AMG232 and RT were targeting S and M-phase cells and promoting G0/G1 cycle arrest (Supplementary Figure 8B). While RT was successfully diminishing mitotic cells across the whole PDCL cohort, this population remained intact upon AMG232 treatment in the *TP53*^MUT^ models. Interestingly, in the *TP53*^MUT^ group, we observed a relative enrichment of cells in S-phase upon RT. This prolonged S-phase might indicate that S-phase checkpoint and DNA damage repair were actively taking place [28] (Supplementary Figure 8A). To understand whether p21 was a mediator of cell cycle arrest, we mapped the cell cycle phases in therapy responsive and unresponsive cells in AMG232- and RT-treated samples (Supplementary Figure 8C & D). In *TP53*^WT^ models, the AMG232-responsive cells (named p21-High) were arrested in G0/G1 phase, while this population in *TP53*^MUT^ cell lines maintained an active cell cycle. RT had a comparable effect on both *TP53*^WT^ and *TP53*^MUT^ PDCLs. Interestingly, upon RT we could identify substantial amounts of p21-Low cell populations in the *TP53*^WT^ PDCL group which were retaining proliferative capacity (Supplementary Figure 8C & D). G0/G1 arrest was only recorded in p21-High populations, independent of the pATM or pH2AX expression status.

### Functional data more accurately capture tumor-specific vulnerabilities than baseline features

Next, we interrogated which of the various drug-perturbation effects from either treatment, including (i) the changes in drug-dependent molecular signatures and/or (ii) cell cycle (Supplementary Figure 9) or (iii) the baseline features of only control samples (Supplementary Figure 10 & 11), would be the most optimal in predicting the cytotoxic outcome through PLI. Using a general linear model of the single and combined drug-specific and cell cycle pseudotime values (for either AMG232 or RT), we observed that the predictive capacity of the changes in drug-induced molecular signatures alone outperformed the readout of the cell cycle alone (Figure 3G & H). Additionally, the combination of both cell cycle and drug-specific readouts did not further improve the PLI prediction of the drug-related molecular signature (Figure 3G & H – Functional response). Furthermore, the therapy perturbation signatures in context of functional readouts outperformed the static measurements in PLI correlation, supporting our hypothesis that functional readouts are more predictable of treatment tolerability of tumor cells (Figure 3 G & H - Static analysis).

### The PROSPERO assay predicts treatment efficacy in fresh brain tumor samples

To evaluate the feasibility of functional testing on fresh GBM samples with the PROSPERO assay, we treated 39 *ex vivo* tumor samples from 34 glioblastoma and high-grade glioma patients with AMG232 and vehicle. Only a subset of samples could also be exposed to RT because of insufficient tumor material. Patient demographics, diagnosis and disease onset (new/recurrent tumor), tumor region (core/invasion) are summarized in Table 1. The treatment conditions that were adapted and optimized in the PDCL pool were applied on the GBM samples (Materials and Methods; Supplementary Table 2). After cell deconvolution into malignant (SOX2+) and non-malignant populations (CD45+/CD3+/CD68+) (Supplementary Figure 12A & B), the relative composition of each sample was determined and ranked. Overall, this analysis showed that myeloid- and T-cells were preserved after acute, functional testing with the PROSPERO assay, even though the general composition of each sample was highly variable (Supplementary Figure 12C). Once cells were deconvoluted according to the applied treatment (control, AMG232 or RT), the drug response analysis and the definition of responsive vs non-responsive populations (see Materials and Methods, Supplementary Figure 13) took two strategic directions: (i) Using the literature-guided drug response signatures and (ii) unbiased, pseudotime analysis and prediction of cellular outcome using the PLI metric (Figure 4A).

**Fig 4.**
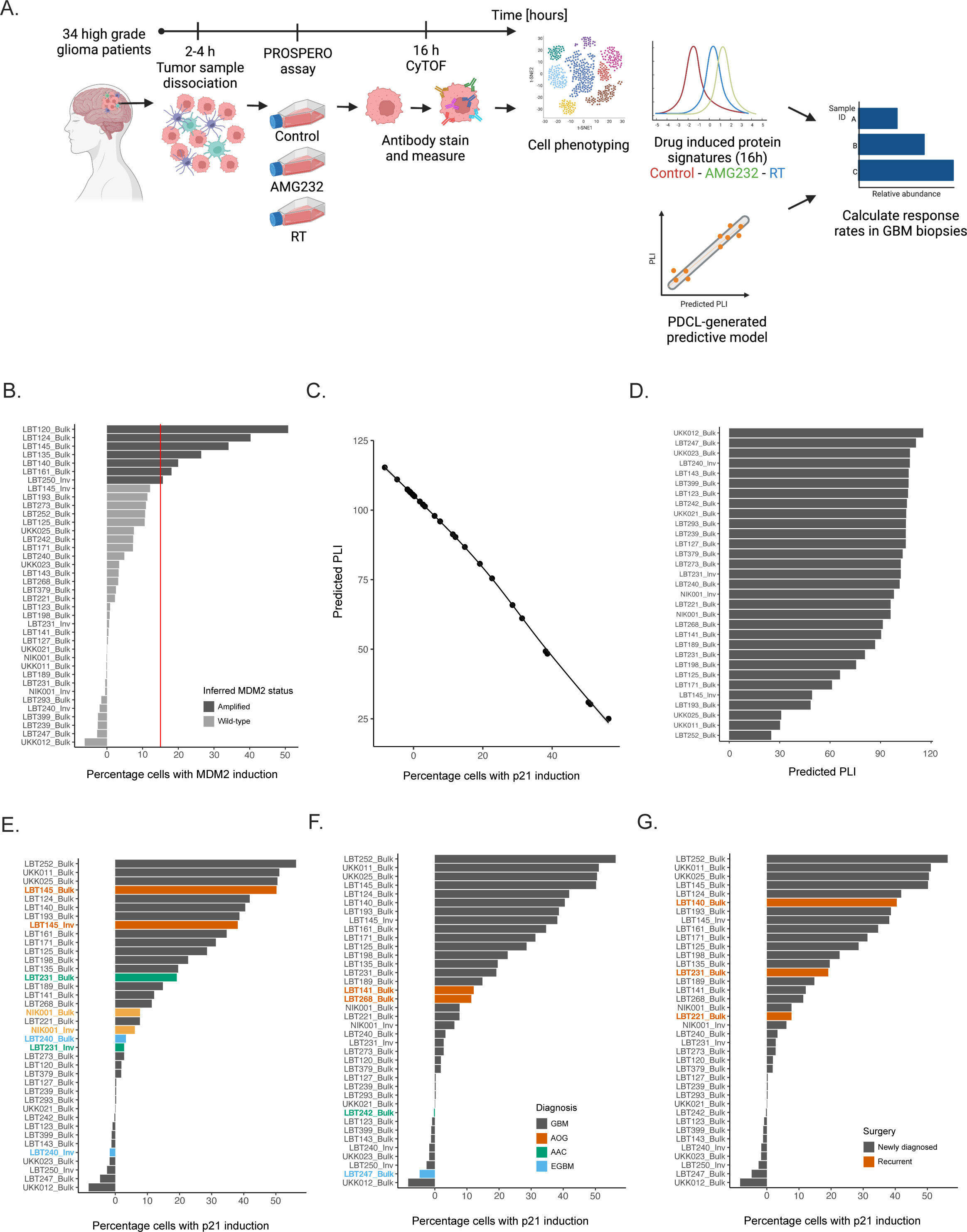
Applying PROSPERO on freshly resected GBM biopsies. (a) Schematic of PROSPERO’s workflow. (b) Barplot ranking based on MDM2 levels. The cutoff level was assessed in the PDCLs (vertical dashed line) and every sample exceeding the cutoff value is considered as an *MDM2*^AMP^. (c) A pseudotime correlation between p21 evels and predicted PLI. (d) Ranking of patients’ samples based on predicted PLI ranging from least (top) to most sensitive (bottom). (e-g) Ranking of patients’ samples based on proportions of responsive cells and colored by: (e) tumor region (core vs invasion); (f) diagnosis; (g) surgery-newly vs recurrent tumor

**Table 1.** Patient demographics from which clinical samples were derived. Characterization of GBM specimens. *^a^*Newly-Diagnosed; *^b^*Recurrent tumor; *^c^*Anaplastic Oligodendroglioma, Gr III; *^d^*Anaplastic astrocytoma, Gr III; *^e^*Epitheloid GBM, Gr IV.

| Surgery | Diagnosis | Sample_ID | Region | TP53-IHC pathology | TP53 Bulk DNaseq | Age | Gender |
| --- | --- | --- | --- | --- | --- | --- | --- |
| ND<sup>a</sup> | GBM | LBT120 | Bulk | MUT | WT | 53 | M |
| ND<sup>a</sup> | GBM | LBT123 | Bulk | MUT | NA | 83 | M |
| ND<sup>a</sup> | GBM | LBT125 | Bulk | MUT | WT | 68 | M |
| REC<sup>b</sup> | GBM | LBT127 | Bulk | MUT | R249S/WT | 51 | M |
| ND<sup>a</sup> | GBM | LBT135 | Bulk | MUT | P64L | 72 | M |
| REC<sup>b</sup> | GBM | LBT140 | Bulk | MUT | WT | 41 | M |
| ND<sup>a</sup> | AOG<sup>c</sup> | LBT141 | Bulk | WT | WT | 73 | M |
| ND<sup>a</sup> | GBM | LBT143 | Bulk | MUT | K132E/WT | 75 | M |
| ND<sup>a</sup> | GBM | LBT145 | Bulk | MUT | WT | 46 | F |
| ND<sup>a</sup> | GBM | LBT145 | Invasion | MUT | WT | 46 | F |
| ND<sup>a</sup> | GBM | LBT161 | Bulk | MUT | WT | 63 | M |
| ND<sup>a</sup> | GBM | LBT171 | Bulk | MUT | NA | 69 | F |
| ND<sup>a</sup> | GBM | LBT189 | Bulk | MUT | NA | 44 | F |
| ND<sup>a</sup> | GBM | LBT193 | Bulk | MUT | NA | 66 | F |
| ND<sup>a</sup> | GBM | LBT198 | Bulk | MUT | NA | 70 | M |
| ND<sup>a</sup> | GBM | LBT198 | Invasion | MUT | NA | 70 | M |
| REC<sup>b</sup> | GBM | LBT221 | Bulk | MUT | NA | 40 | M |
| REC<sup>b</sup> | GBM | LBT231 | Bulk | MUT | S9N/P64L | 50 | M |
| REC<sup>b</sup> | GBM | LBT231 | Invasion | MUT | WT | 50 | M |
| ND<sup>a</sup> | GBM | LBT239 | Bulk | MUT | WT | 60 | M |
| ND<sup>a</sup> | GBM | LBT240 | Bulk | MUT | R282W | 57 | F |
| ND<sup>a</sup> | GBM | LBT240 | Invasion | MUT | NA | 57 | F |
| ND<sup>a</sup> | AAC<sup>d</sup> | LBT242 | Bulk | MUT | NA | 21 | F |
| ND<sup>a</sup> | EGBM<sup>e</sup> | LBT247 | Bulk | MUT | G199E | 30 | F |
| ND<sup>a</sup> | GBM | LBT250 | Invasion | MUT | WT | 51 | M |
| ND<sup>a</sup> | GBM | LBT252 | Bulk | MUT | WT | 45 | M |
| ND<sup>a</sup> | AOG | LBT268 | Bulk | WT | WT | 27 | M |
| ND<sup>a</sup> | GBM | LBT273 | Bulk | WT | WT | 35 | M |
| ND<sup>a</sup> | EGBM | LBT293 | Bulk | MUT | R282W | 35 | M |
| ND<sup>a</sup> | GBM | LBT379 | Bulk | MUT | 470ins12bp | 69 | M |
| ND<sup>a</sup> | GBM | LBT399 | Bulk | MUT | WT | 69 | M |
| ND<sup>a</sup> | GBM | NIK001 | Bulk | WT | NA | 65 | M |
| ND<sup>a</sup> | GBM | NIK001 | Invasion | WT | NA | 65 | M |
| ND<sup>a</sup> | GBM | UKK011 | Bulk | WT | WT | 52 | F |
| ND<sup>a</sup> | GBM | UKK012 | Bulk | WT | NA | NA | NA |
| ND<sup>a</sup> | GBM | UKK021 | Bulk | MUT | C176W | 44 | M |
| ND<sup>a</sup> | GBM | LBT124 | Bulk | NA | WT | 39 | M |
| ND<sup>a</sup> | GBM | UKK023 | Bulk | MUT | NA | NA | NA |
| ND<sup>a</sup> | GBM | UKK025 | Bulk | WT | NA | NA | NA |

The first strategy confirmed that, in general, the response rates upon AMG232 and RT correspond to the *TP53* mutational status (Supplementary Figure 14). Also, samples from the invasive margin of the tumor were less responsive to therapeutic insults (Supplementary Figure 14B & D).

In the context of AMG232 treatment, the second analysis pipeline followed the PDCL-trained model of *MDM2*^AMP^ (Figure 4B) sample stratification and PLI prediction based on the responsiveness scores in the biopsy samples (Supplementary Figure 15A & B). Thus, predicted PLI values were calculated for the biopsy samples (Figure 4C) and ranked for AMG232 (Figure 4D) and RT (Supplementary Figure 15C).

**Fig 5.**
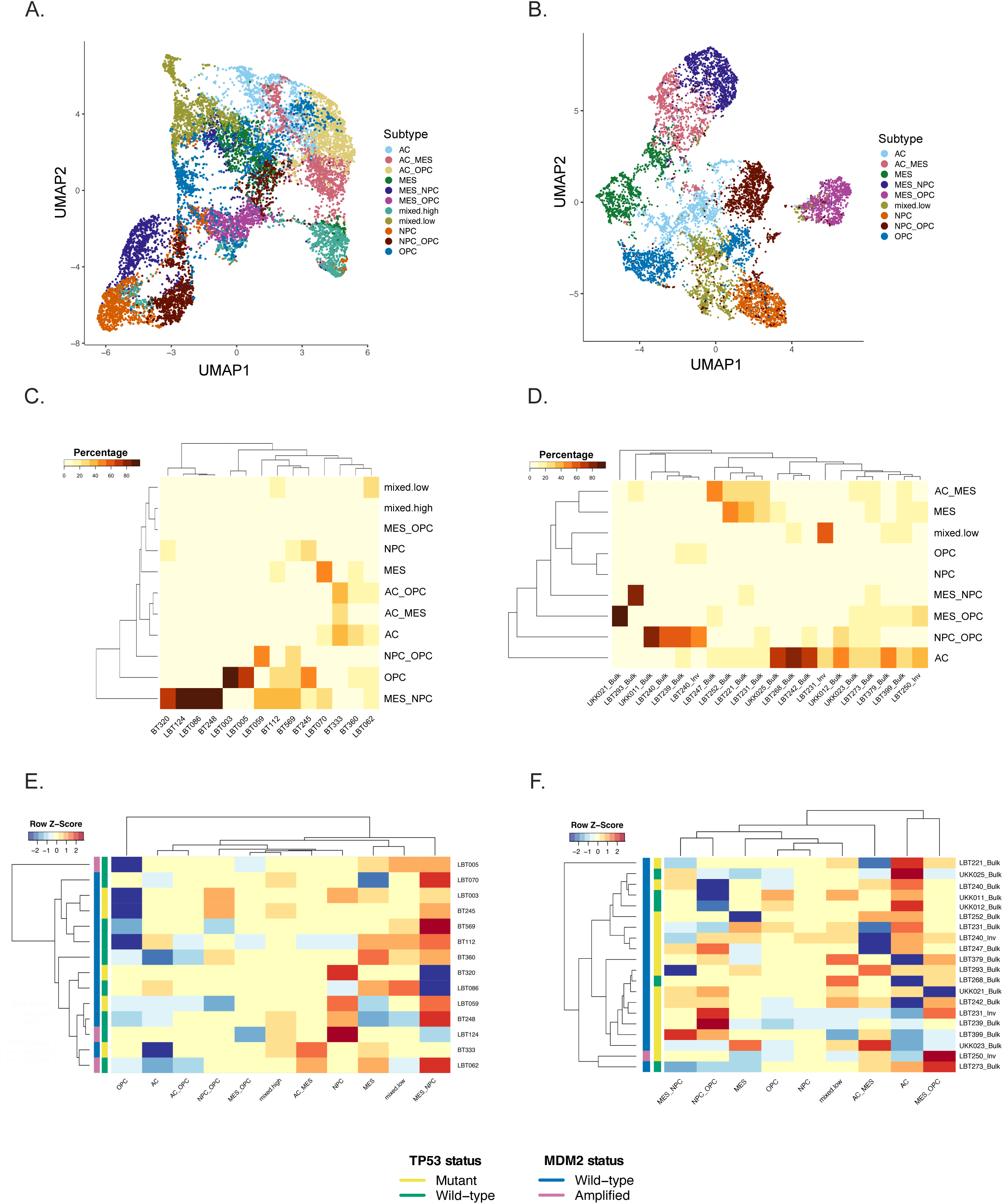
Baseline heterogeneity and therapy-induced plasticity upon AMG232 treatment. UMAP visualization of tumor cells in (a) PDCLs and (b) biopsies (colored by phenotype). Heatmap of proportion of malignant cells assigned to each phenotype across control-treated samples from (c) PDCLs and (d) biopsies. Therapy-induced phenotypic shifts in (e) PDCLs and (f) biopsies

Again, with the predictive model, we observed that samples derived from the invasive tumor margin tended to be less responsive than the corresponding bulk/core samples (Figure 4E). While tumor cells from two astrocytic oligodendrogliomas showed relatively overlapping amounts of responsive cells (Figure 4F), the ability of newly-diagnosed and recurrent GBM samples to respond to therapy was more heterogeneous (Figure 4G). Once again, upon exposure to RT, a variable number of cells were able to demonstrate a proper molecular drug-response, even though this ability generally did not correlate to either genomic/IHC-based *TP*53 status (Table 1).

### The PROSPERO assay quantifies cellular GBM heterogeneity

To determine the ability of PROSPERO to detect GBM heterogeneity at baseline and upon therapy exposure, we assessed the expression of canonical markers associated with the RNA-based classification, which conveniently groups glioma stem-like cells into four major cellular states, including astrocyte-like (AC), oligodendrocyte precursor-like (OPC), neural precursor-like (NPC) and mesenchymal-like (MES) tumor cell states [1] (Supplementary Table 3). To identify subpopulations, unsupervised clustering was performed on both datasets (Supplementary Figure 17A). Then, cells were deconvoluted based on treatment conditions. Subsampled UMAP embeddings of control cells from both datasets are presented in Figure 5A & B; Supplementary Figure 18. The examination of control and treated samples showed that all four states and the transitional lineages between them were well conserved in both datasets. In baseline, the cell lines were mostly represented by mesenchymal-like and neuronal-like progenitors. Only two models (LBT003 and LBT005) were populated particularly by oligodendrocyte-like precursor cells (Figure 5C). In contrast, half of the biopsy specimens contained only astrocyte-like (GFAP+) cells, while the other ten samples contained either mixes of mesenchymal cells with the other three phenotypes or proneural cells (NPC/OPC) (Figure 5D).

Furthermore, we wanted to inspect the proliferation capacity of the phenotypic subclasses in baseline samples (as described in Material and Methods) [22] (Supplementary Figures 7 & 19). In the PDCL pool, NPC and NPC/OPC mixtures seemed to retain a higher proliferative potential (lowest percentage of G0/G1 cells and highest number of S-phase cells) [22] (Supplementary Figure 19C). In contrast, patients’ samples expressed larger percentage of cells in cell cycle arrest (G0/G1), emphasizing the inability of standard-of-care treatment using TMZ/RT to target these populations through the cell cycle machinery (Supplementary Figure 19B).

### Therapy-induced plasticity as response to therapeutic intervention across PDCLs and GBM biopsies

In general, the PDCLs exhibit quintessential proneural (NPC/OPC)-to-mesenchymal transitions upon AMG232 treatment (Figure 5E; Supplementary Figure 20A). The depletion of OPC-like (OLIG2+/PDGFRa+) populations could be mainly explained by the upregulation of *TP*53 (in *TP53*^WT^ models) and the concurrent downregulation of PDGFRa, caused by p73 activation [32] (Supplementary Figure 3B; Supplementary Figure 8B). When looking at percentages of responsive cells, the enrichment of MES-like phenotypes in the PDLCs is presented only in the *TP53*^WT^ group, similarly as in the biopsies (Supplementary Figure 20A & B). However, the effect of AMG232 on the phenotypic transition in the patients’ samples showed a bidirectional pattern, often exchanging proneural lineages (NPC/OPC) with astro-mesenchymal (AC/MES) states or vice versa (Figure 5F). Upon RT, the neural-like progenitors were the responsive cells in our PDCL dataset (Supplementary Figure 20C). We observed a substantial downregulation of OPC populations (mainly in the *TP53*^WT^ group) and a phenotypic transition commonly towards mesenchymal phenotype (Supplementary Figure 20E). In the small number of RT treated biopsy samples, MES/NPC were the most responsive populations (Supplementary Figure 20D) and the direction of the phenotypic transition is rather patient-dependent (Supplementary Figure 20F).

### *Ex vivo* tumor cells isolated from hetero- and ortho-topical PDX models retain ***in vivo* responsiveness profiles**

The *in vivo* activity of MDM2 inhibitors has previously been investigated in patient-derived xenograft models [19] and showed survival benefits across a number of *TP53*^WT^ and *MDM2*^AMP^ models, including the BT112 cell line used in this study. Here, we intracranially inoculated mice with GFP/luciferase-positive BT112 and treated one group with AMG232 by oral gavage, while the other group received vehicle control. After 16 hours, mice were sacrificed and living tumor tissue was dissociated as describe above (Figure 6A, Supplementary materials). Subsequently, we compared the molecular profiles in tumor cells isolated from the mice that were treated with AMG232 by oral gavage, with tumor cells from control mice that were exposed to AMG232 treatment *ex vivo* using the PROSPERO assay. First, we observed that drug-response signatures between *ex vivo* PROSPERO (PDX01-03) and *in vivo* treated mice (PDX04-07) were very comparable. In addition, when comparing the extent and prevalence of the changes in marker expression in the mouse-derived tumor samples with the original GFP/luciferase-positive BT112, signatures were less pronounced and prevalent in the mouse-derived samples (Figure 6B). This agrees with the discrepancy that AMG232 is able to efficiently kill all BT112 cells when maintained as a PDCL, while the *in vivo* ability of AMG232 to cure the PDX mice was largely confined to delaying disease progression [19]. Overall, this experiment shows that the ex vivo PROSPERO assay correlates with the drug activity that is ongoing *in vivo*. Future clinical trials will have the provide evidence as to whether the insights of PROSPERO are also translatable to identify the most active treatment in GBM patients.

**Fig 6.**
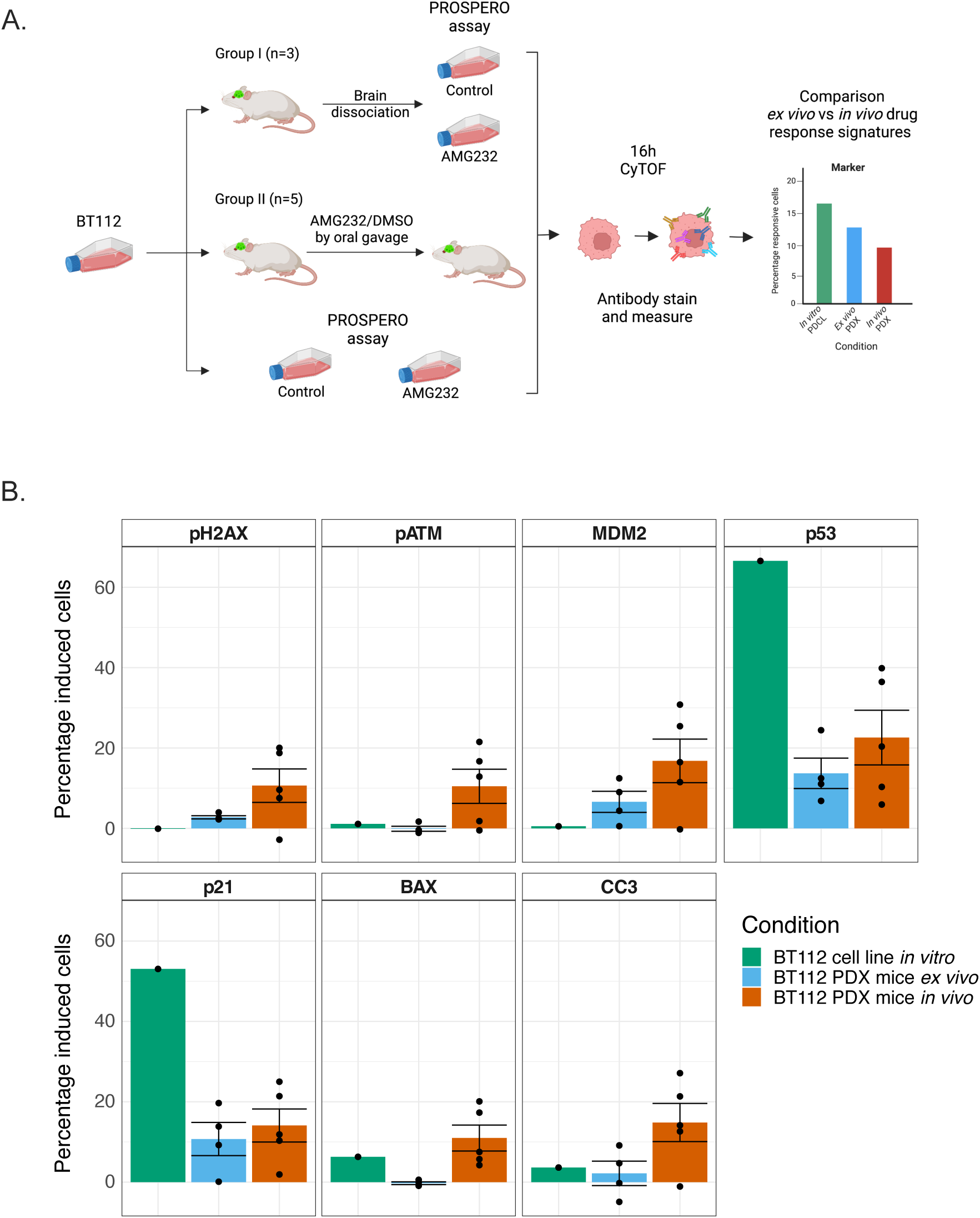
Validation of PROSPERO in PDX model. (a) Schematic diagram of experimental setup. (b) Percentage of cells showing drug-related marker induction (mean ± SD) in BT112 (PDCL), *ex vivo* treated cells with PROSPERO and cells isolated from *in vivo* treated mice

## Discussion

Inspired by the recent success of a functional diagnostics approach introduced in advanced hematological malignances [15, 16], we developed a brain-tumor-focused functional readout to interrogate therapeutic efficacy at single-cell resolution, called the PROSPERO assay.

GBM samples were treated *ex vivo* within few hours post-surgery allowing the assessment of crucial molecular and cellular signatures of drug response within a clinically relevant timeframe (ie 2-3 days post-surgery). Importantly, by applying this strategy to tumor samples derived from distinct tumor regions, glioblastoma and high-grade glioma diagnoses and tumor relapses, we showed the feasibility and convenience of this strategy in providing therapeutic insights at any moment during the treatment scheme of a patient.

By measuring the molecular changes induced by either therapy at single-cell resolution, we revealed patients’ sample-specific response heterogeneity and therapy-induced plasticity. These two observations were found to only poorly correlate to static parameters such as, bulk genetic or IHC - *TP53* mutational status [33] (Table 1), suggesting that baseline *TP53* mutational assessment may be of insufficient precision to make a proper estimate of therapy responsiveness. IHC staining of p53 and corresponding H&E of few biopsy samples are showing heterogeneous p53 expression (Supplementary Figure 16), supporting our hypothesis that functional testing may deliver a more accurate prediction of the intrinsic capabilities of tumor cells to respond to a given therapy, in comparison to standard techniques.

Collectively, in only one third of the biopsies, AMG232 was able to induce drug responses in a sufficiently large population of tumor cells. A higher degree of sensitivity was noted in *MDM2^AMP^* samples [19], which is typically only a small subgroup of GBM patients (10% - 15%) [34]. RT demonstrated to be less capable to achieve extensive drug responses, even though it was effective at targeting M-phase cells. Stable p53/p21 upregulation is necessary for the activation of cellular senescence, but functional tumor suppressor CDKN2A/p16 is required to maintain the senescent state [35]. Namely, *CDKN2A* deletion is one of the major GBM’s hallmarks [4], which is also presented in both *TP53* pools (Supplementary Figure 3B) aiding tumor cells to circumvent therapy-induced senescence. This might further explain the intrinsic inability of the analyzed cells to present a genuine drug response.

Moreover, intrinsically resistant cell populations from the tumor edge raise an important point, as these populations are too risky to surgically remove and remain infiltrating into the surrounding brain tissue, highlighting the need for rapid, precision tools capable of identifying effective therapeutic agents.

Here, we now show that, already within hours following drug perturbation, tumor cells can change their phenotypes. With PROSPERO we were able to confirm that AC/OPC lineages are mainly enriched in fresh tumor biopsies, which are considered to be treatment-resistant and responsible for recurrence onset [36, 37]. Furthermore, we captured the diversity in phenotypic transitions between tumor core and invasion samples, whereby infiltrating cells mainly gain MES/OPC lineages. MES precursors are considered to be pro-inflammatory indicators of early treatment response [37, 38]. How the observed plasticity correlates to resistance at later stages still remains an outstanding question, but the differential ability of tumor cells to exhibit measurable plastic behavior adds another functional insight.

A primary challenge of this study is that, we have not yet been able to directly correlate the functional observations from the *ex vivo* treatments to patients’ responses. In order to draw clinically relevant conclusions, a larger cohort of patients/samples will have to be *ex vivo* evaluated with a PROSPERO-like approach, while integrating additional, predictive clinical parameters such as extent-of-resection/tumor size/location, number of applied therapy cycles, and genomic aberrations. Recently, the first clinical trials have been initiated, whereby cytotoxicity assays as functional readouts are intended to be performed on patient-derived models and *ex vivo* treated GBM biopsies [6]. Although PDCLs and biopsies are similar, they are not identical [18]. Model propagation/prolonged culturing might be highly time-, labor- and cost-inefficient, hindering the opportunity to provide functional insights for every patient. In this light, PROSPERO was able to capture the similarity of *ex vivo* vs *in vivo* drug responses in PDX models showing that *ex vivo* functional measurements are able to circumvent these limitations.

To our knowledge, this is the first study in GBM deliberating single-cell protein resolution of therapy responses in tumor cells, while comparing diverse tumor regions, diagnosis and disease onset. Our study sets ground for cost/time efficient functional diagnostic assays, which may complement genetic and pathological measurements and accelerate clinical understanding and management of this disease.

## Acknowledgements

The authors thank Prof. Cynthia Guidos and her team from the Department of Immunology, SickKids, for providing guidance in mass cytometry experiments, Tom Janssens for helping perform mouse studies and Nikolina Dubroja-Lakic, Department of Imaging and Pathology, KU Leuven for acquiring IHC images.

## Statements and Declarations

### Author Contributions

All authors contributed to the study conception and design. Material preparation, data collection and analysis were performed by Dena Panovska, Marleen Derweduwe, Tatjana Verbeke, Asier Antoranz and Pouya Nazari. The first draft of the manuscript was written by Dena Panovska and Frederik De Smet and all authors commented on previous versions of the manuscript. Review and editing were done by Basiel Cole, Dena Panovska and Frederik De Smet. All authors read and approved the final manuscript.

## Data availability

The datasets generated and analysed during the current study are not publicly available at the moment, but are available from the corresponding author upon manuscript publication or on reasonable request.

## Ethics approval

The collection and study on human material was approved by University Hospitals Leuven (Leuven, Belgium; IRB protocol S59804), Europaziekenhuizen (Brussels, Belgium; IRB protocol EC approval 05-02-2018), Jessa Ziekenhuis (Hasselt, Belgium, IRB B243201941451), and AZNikolaas (Sint Niklaas, Belgium; IRB protocol EC18021).

All experimental procedures performed on live animals were reviewed and approved by the KULeuven Animal Ethics Committee under the protocol P211/2018. Experiments were conducted in accordance with the KULeuven animal facility regulations and policies.

## Consent to participate

Written informed consent was obtained from the parents.

## Consent to publish

The authors affirm that human research participants provided informed consent for publication of the images in Supplementary Figure 16.

---

**Supplementary Figure 1.**
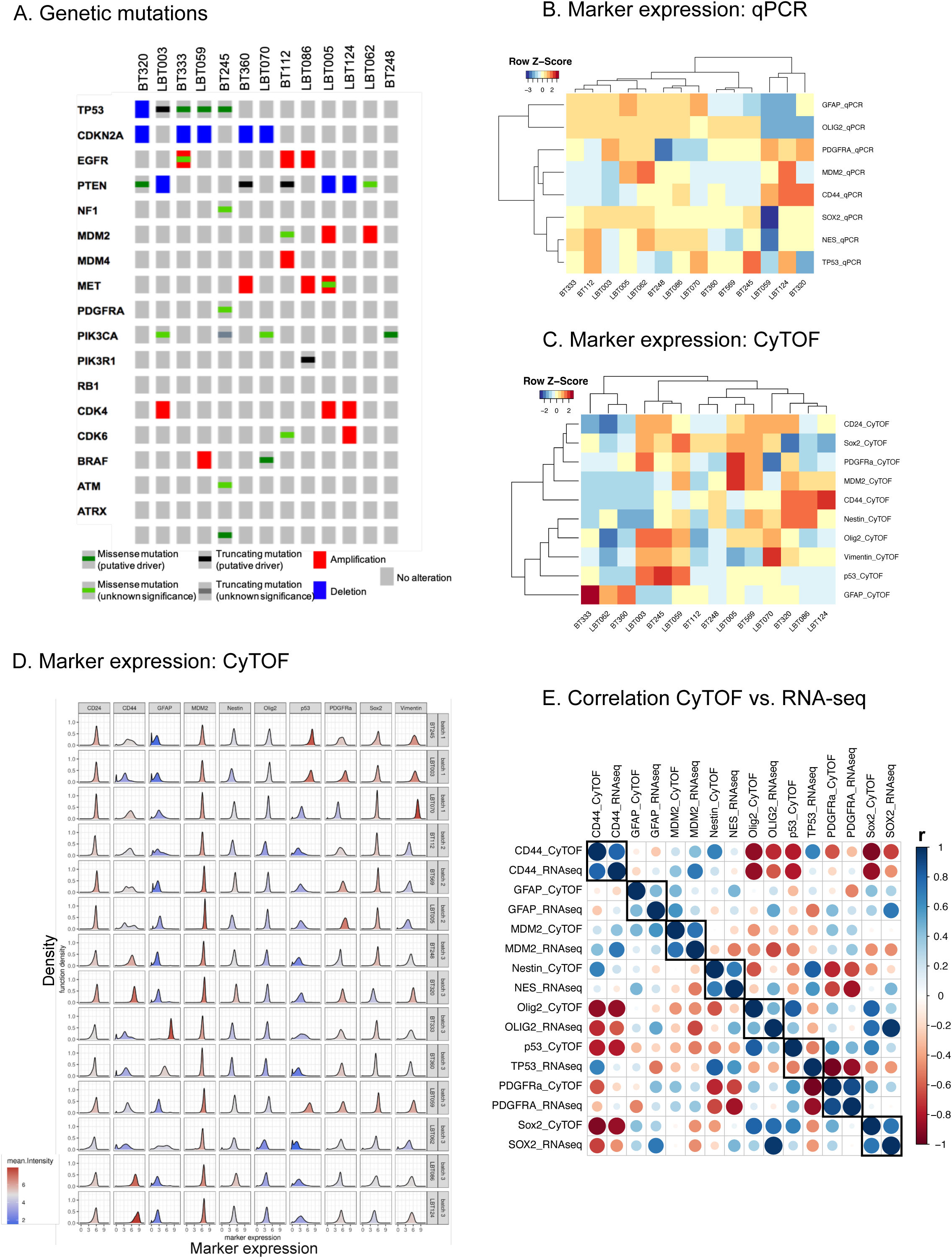

**Supplementary Figure 2.**
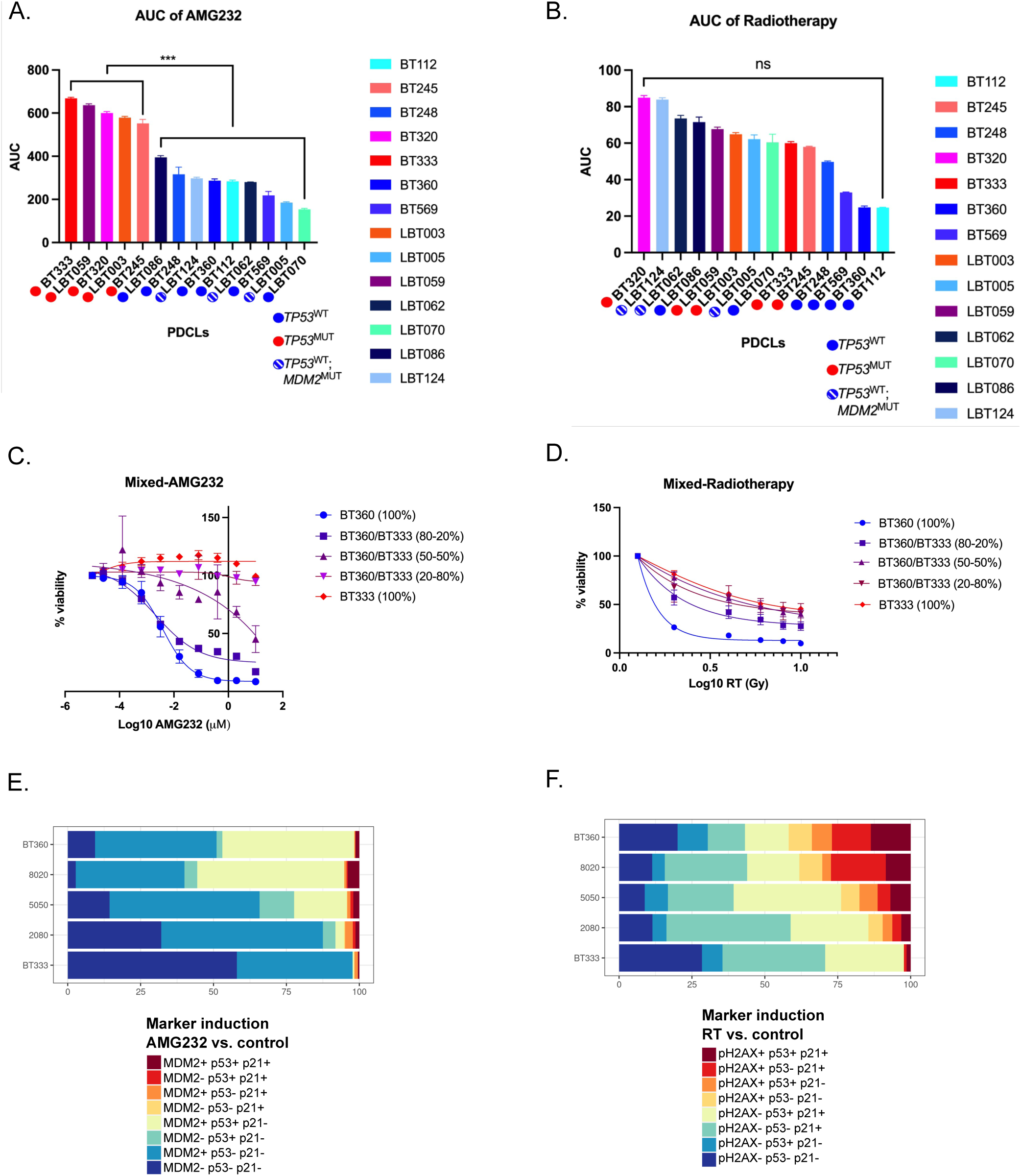

**Supplementary Figure 3.**
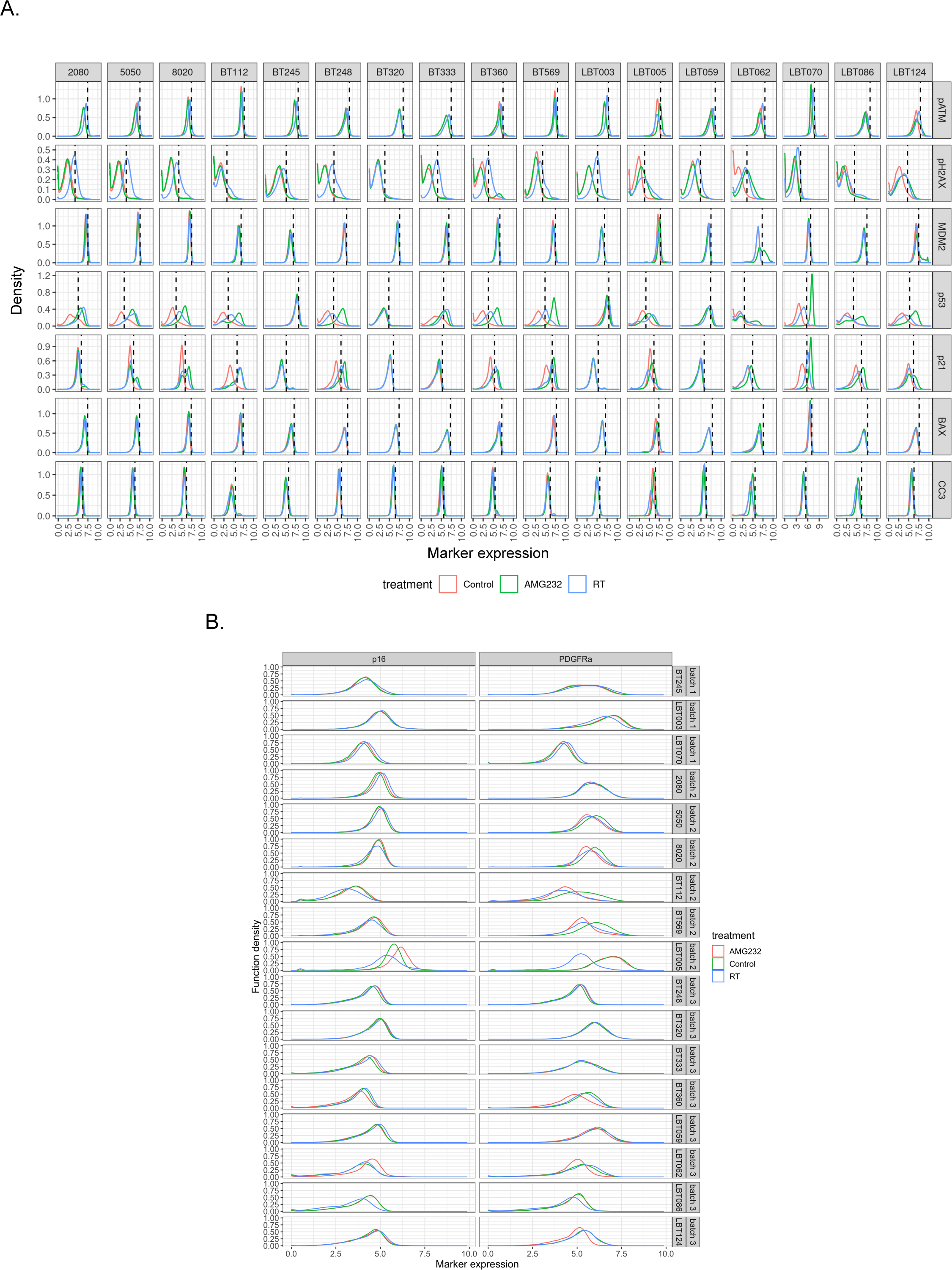

**Supplementary Figure 4.**
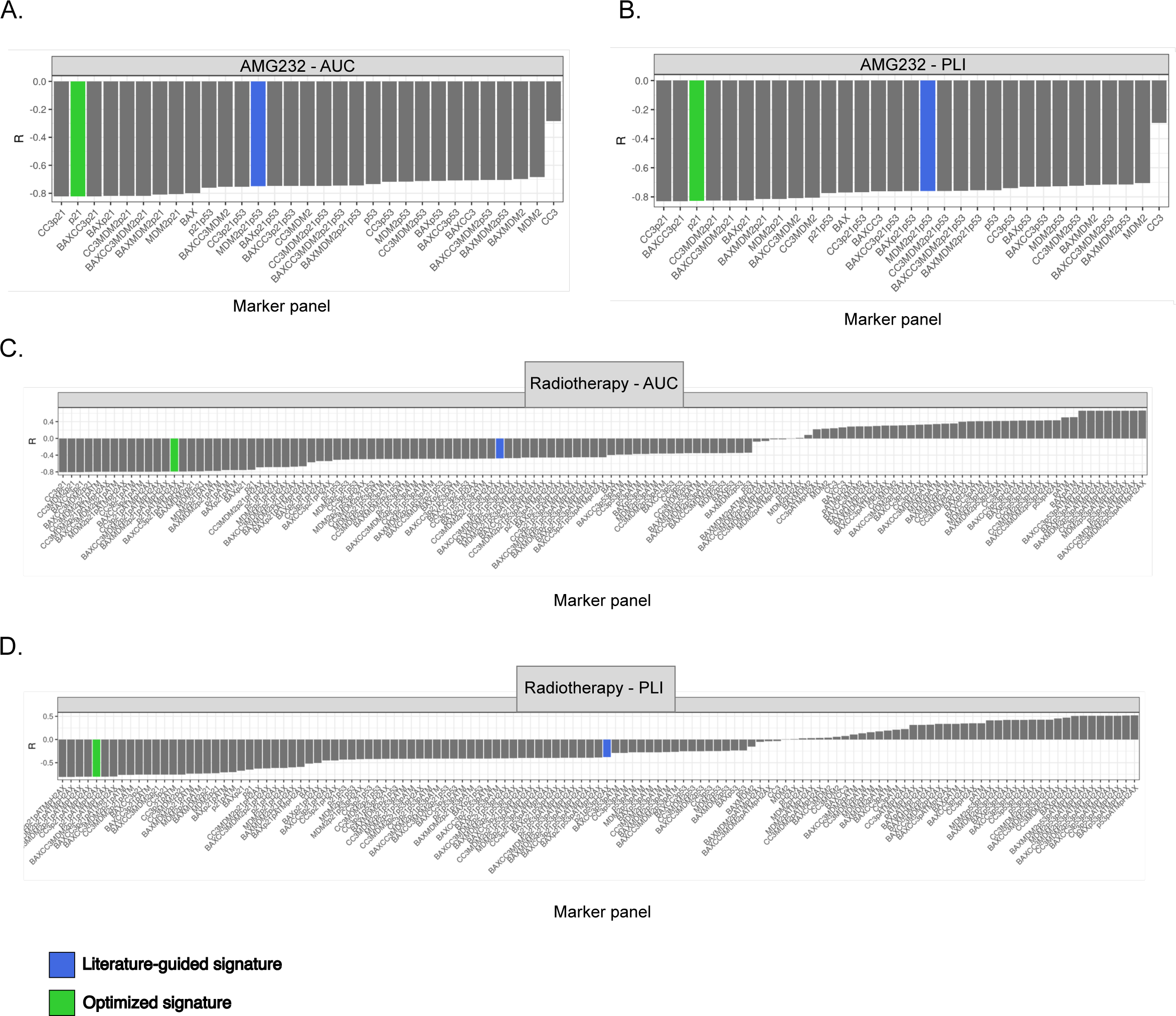

**Supplementary Figure 5.**
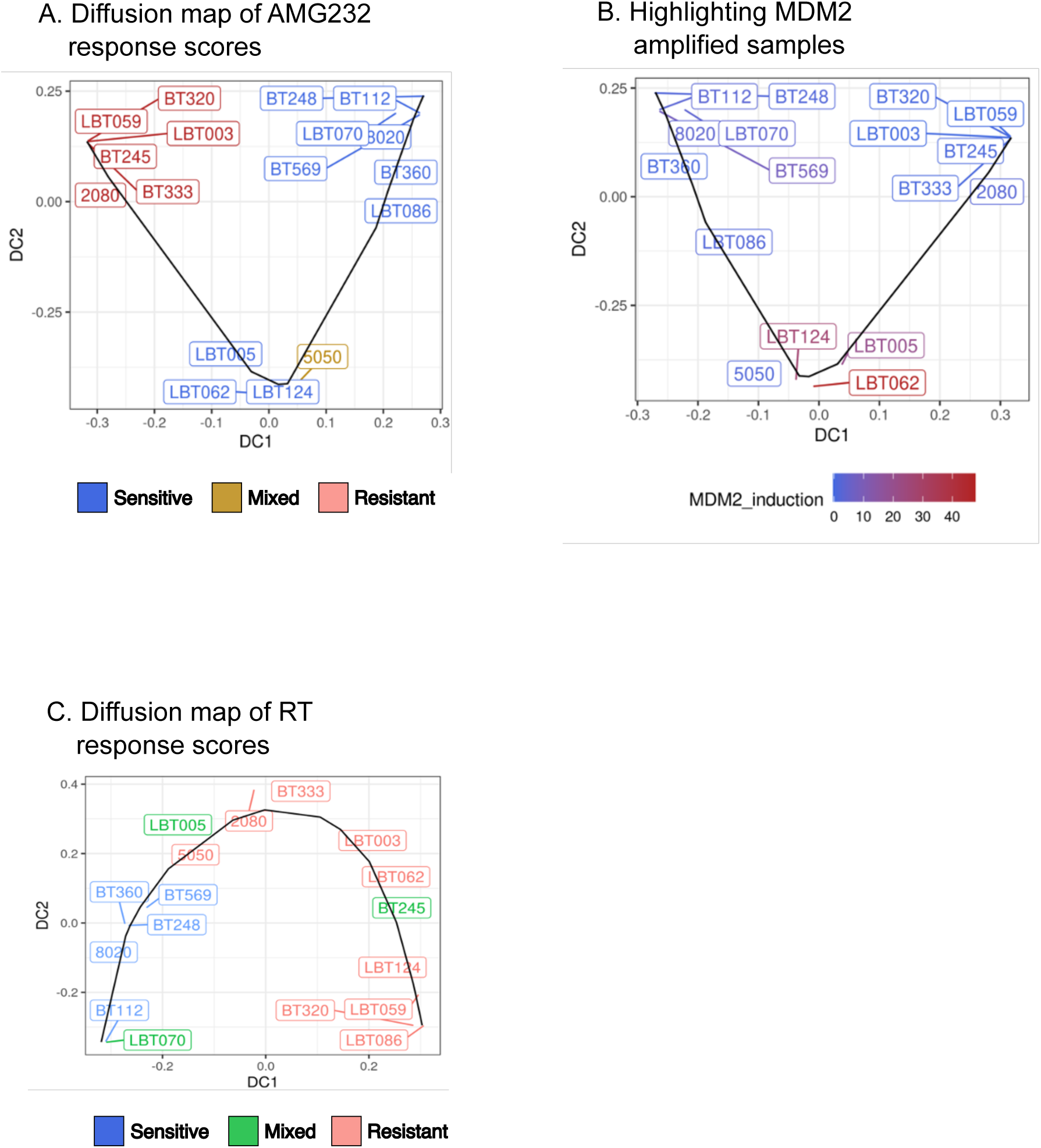

**Supplementary Figure 6.**
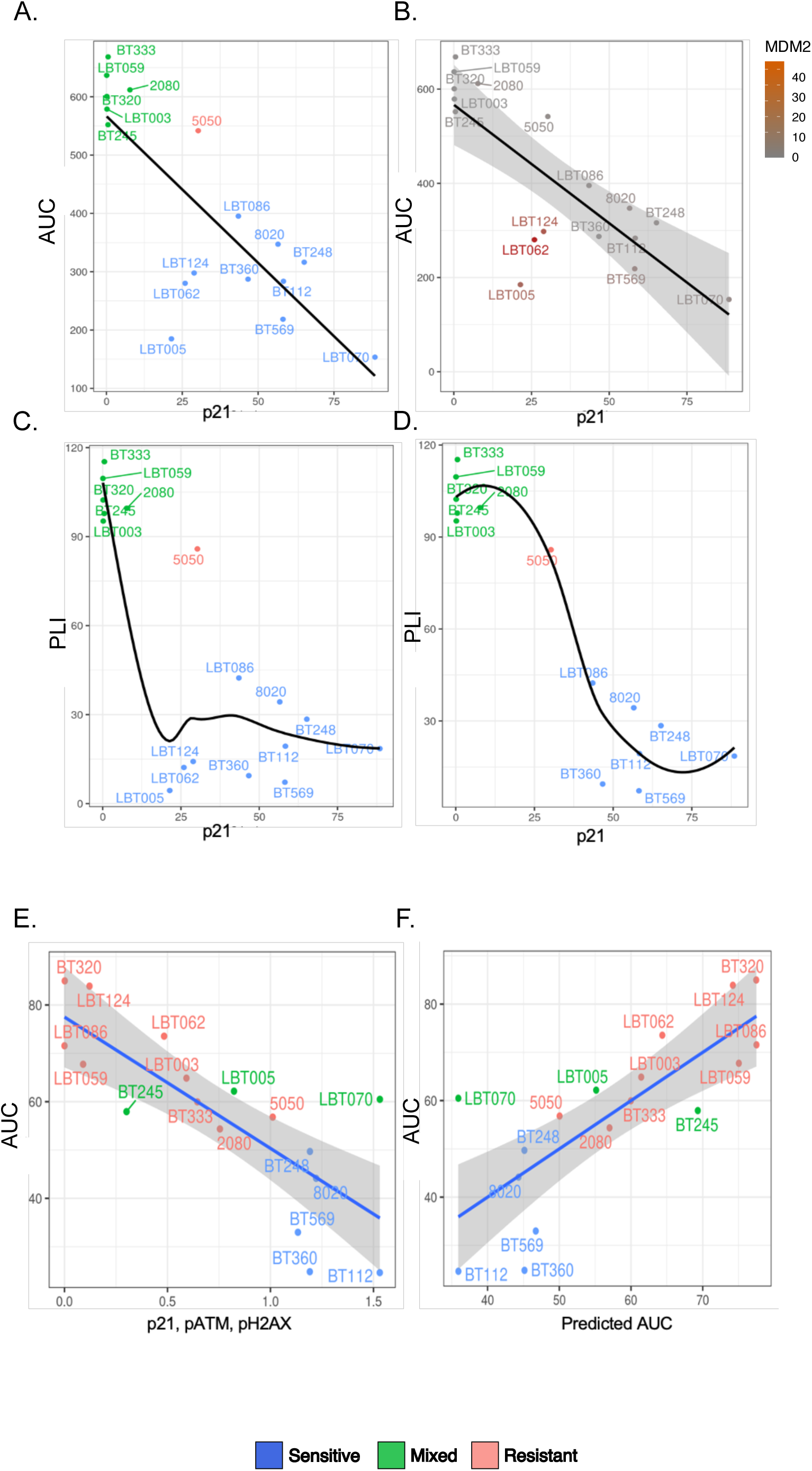

**Supplementary Figure 7.**
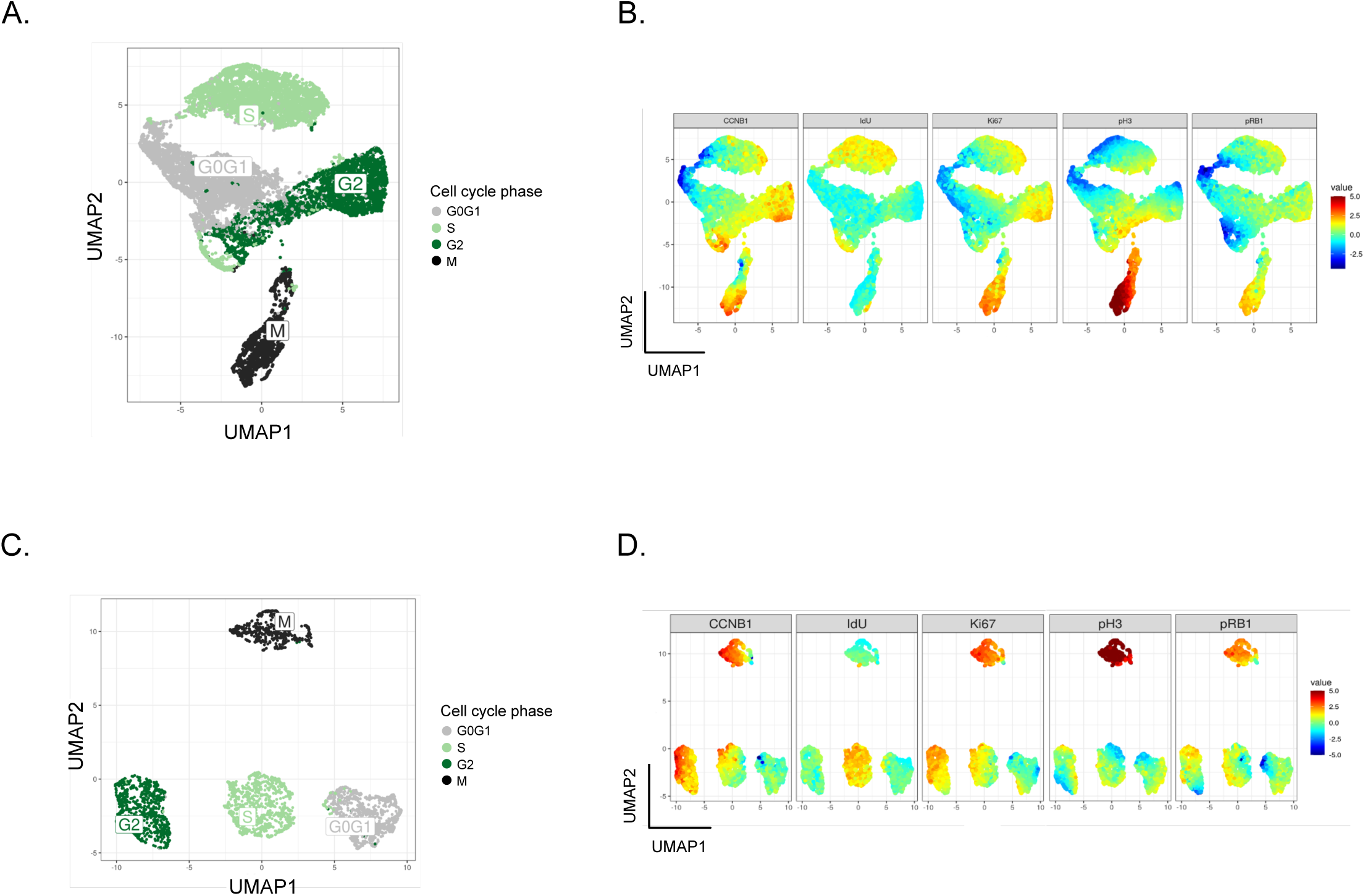

**Supplementary Figure 8.**
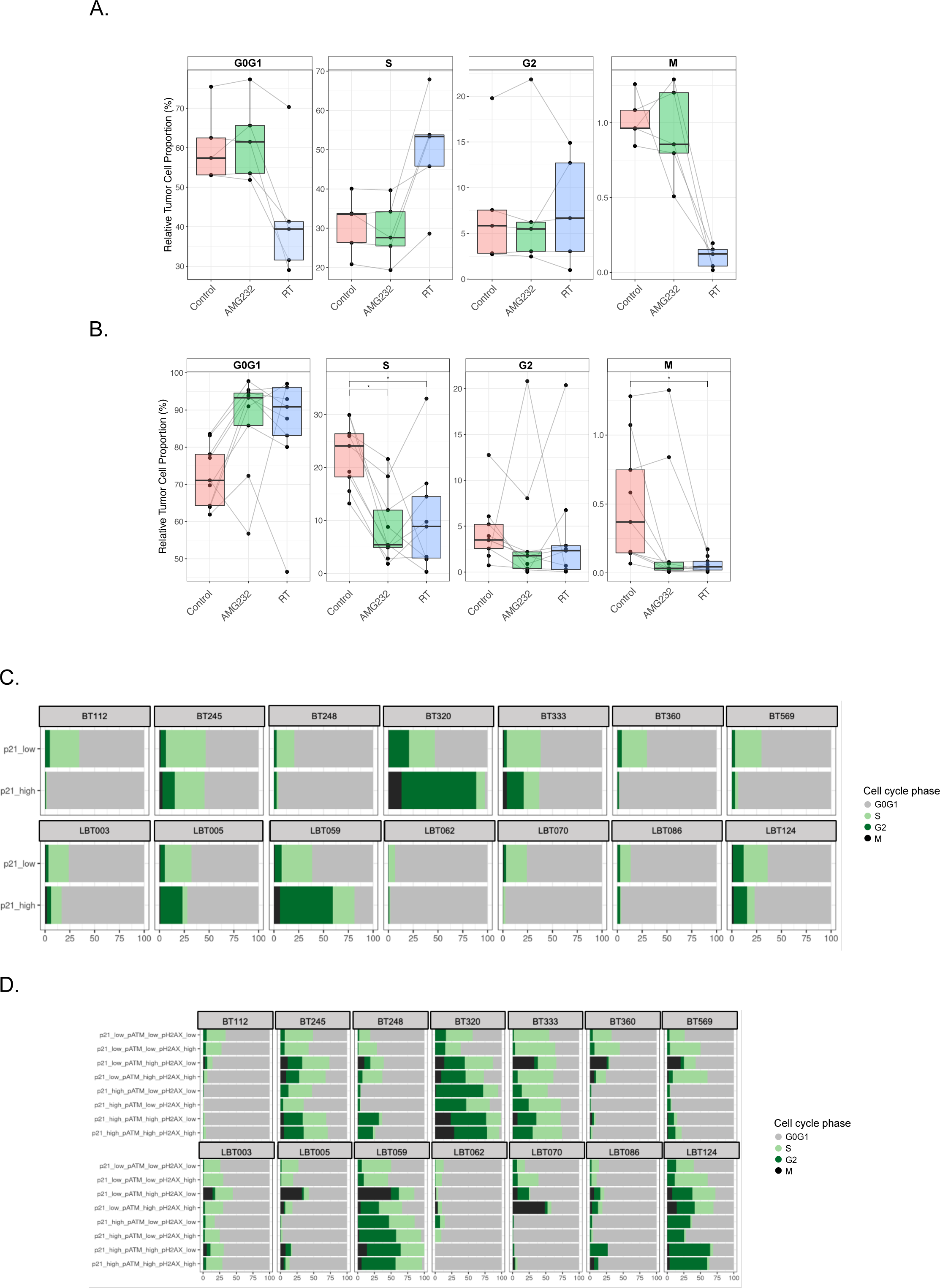

**Supplementary Figure 9.**
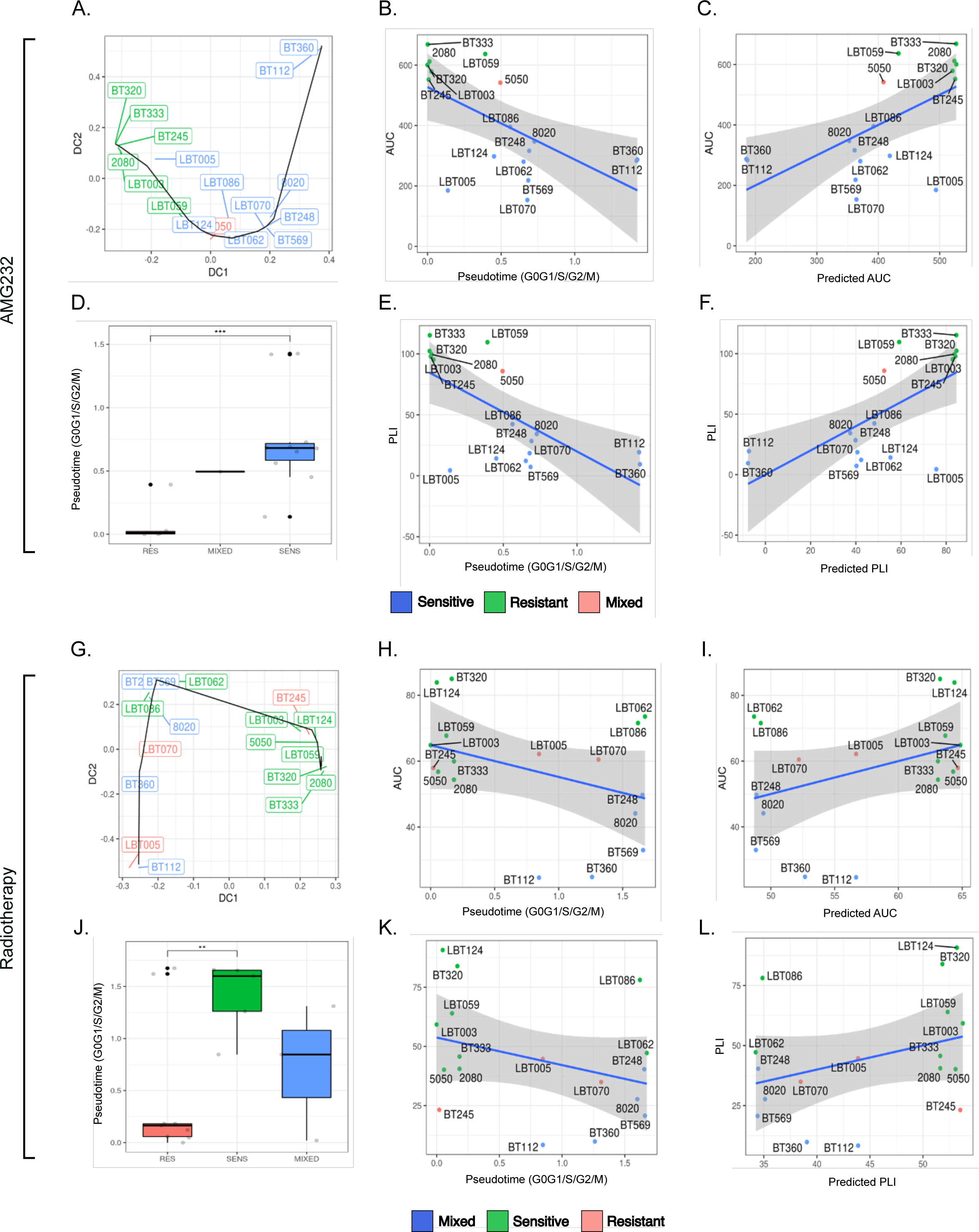

**Supplementary Figure 10.**
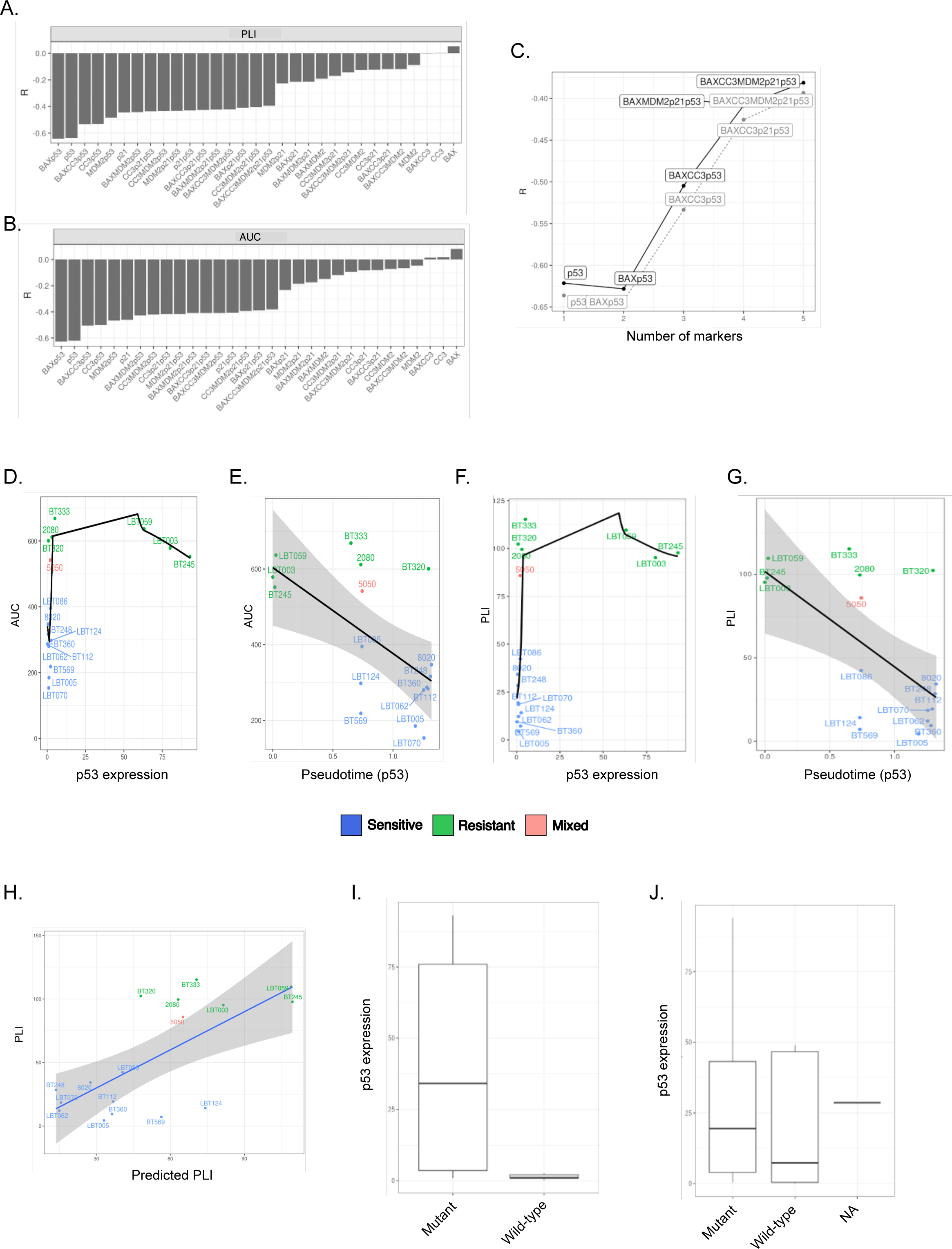

**Supplementary Figure 11.**
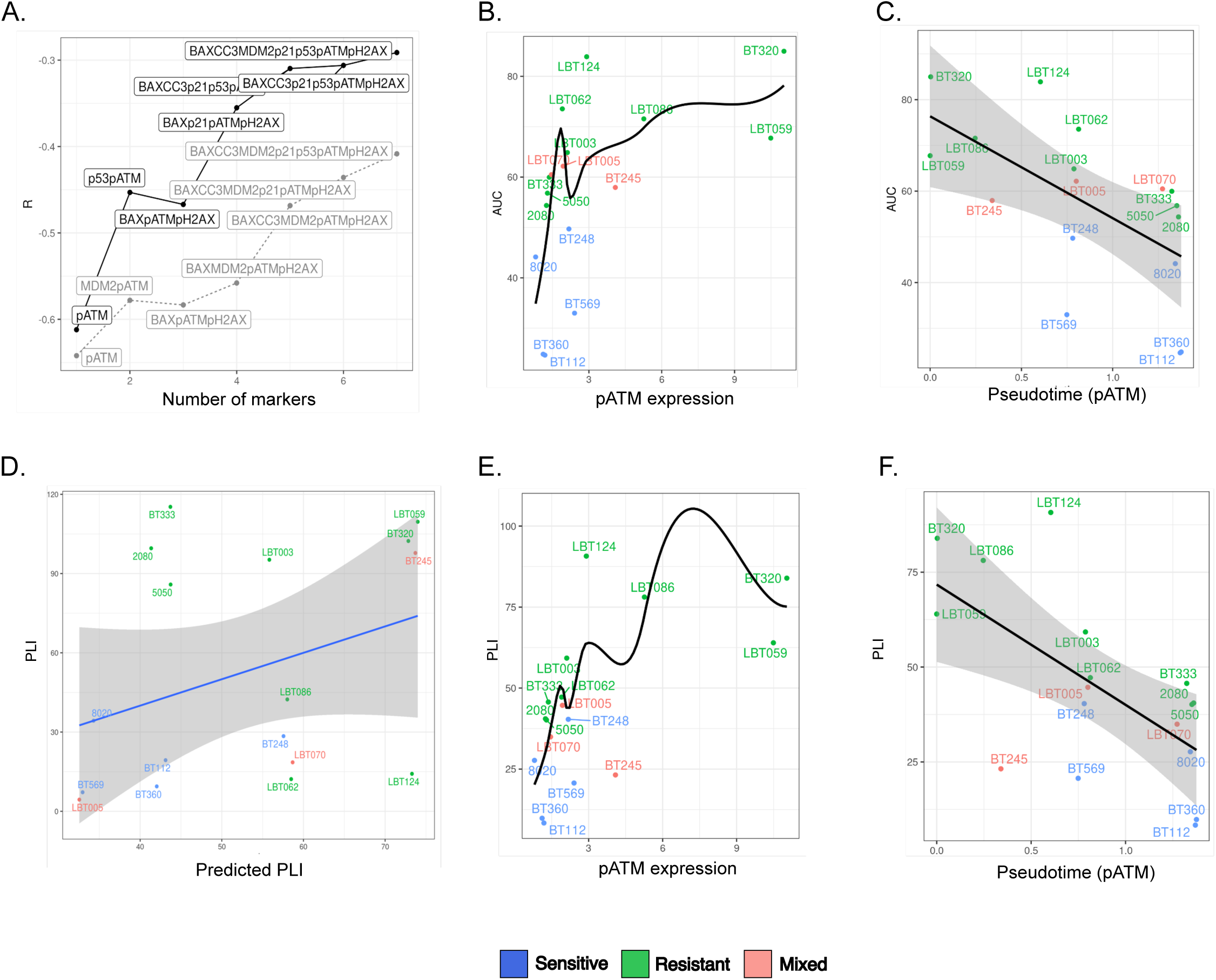

**Supplementary Figure 12.**
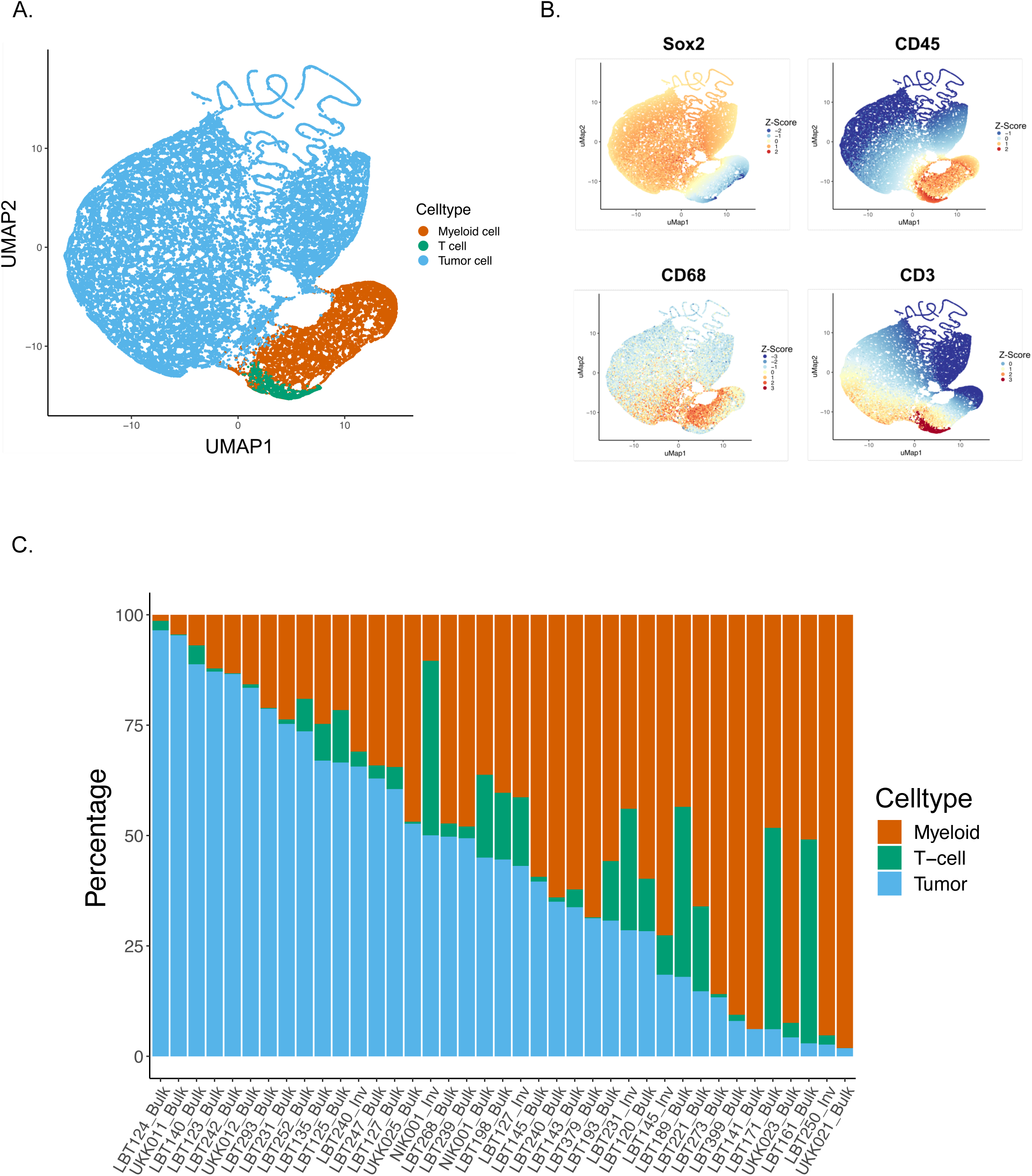

**Supplementary Figure 13.**
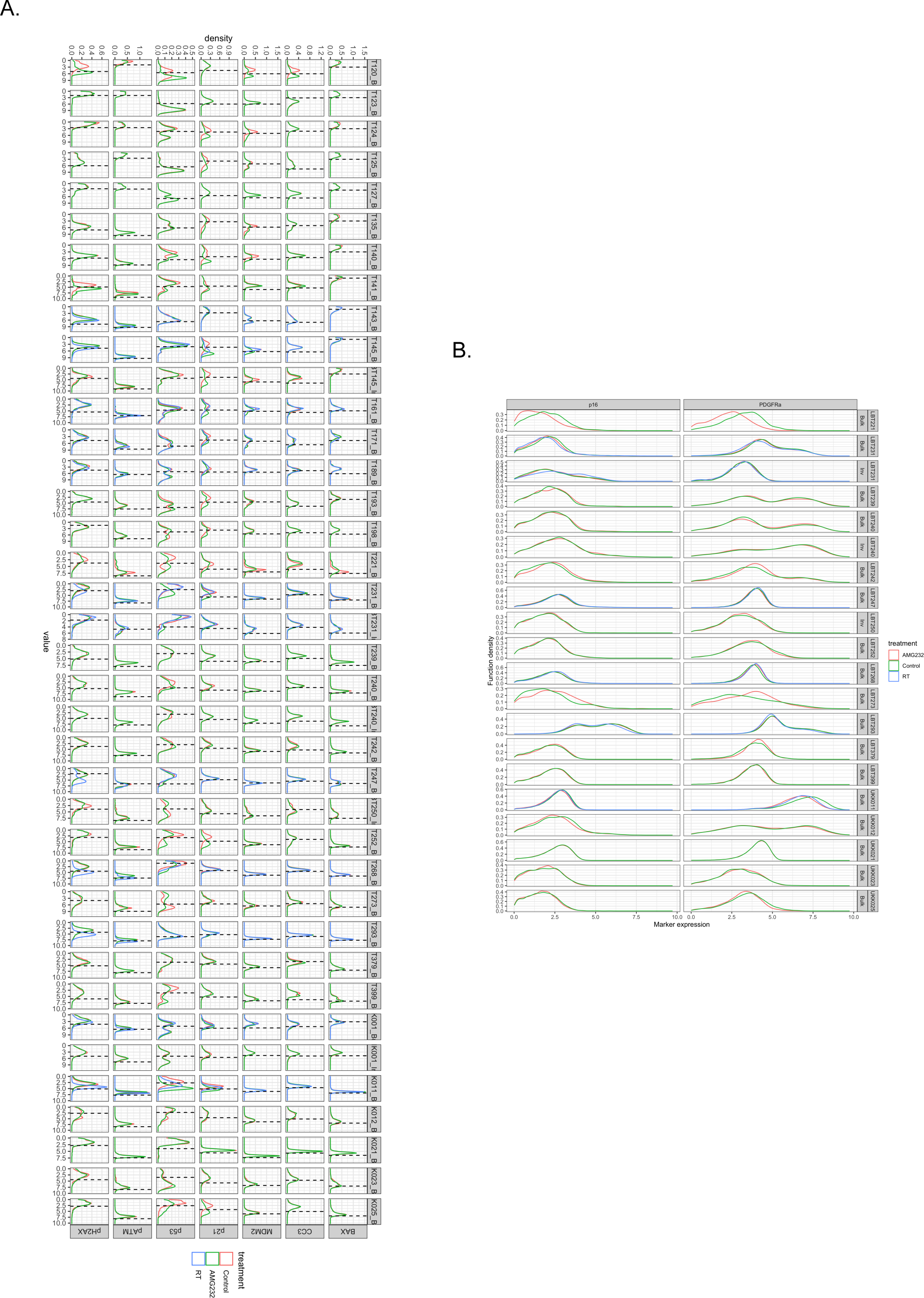

**Supplementary Figure 14.**
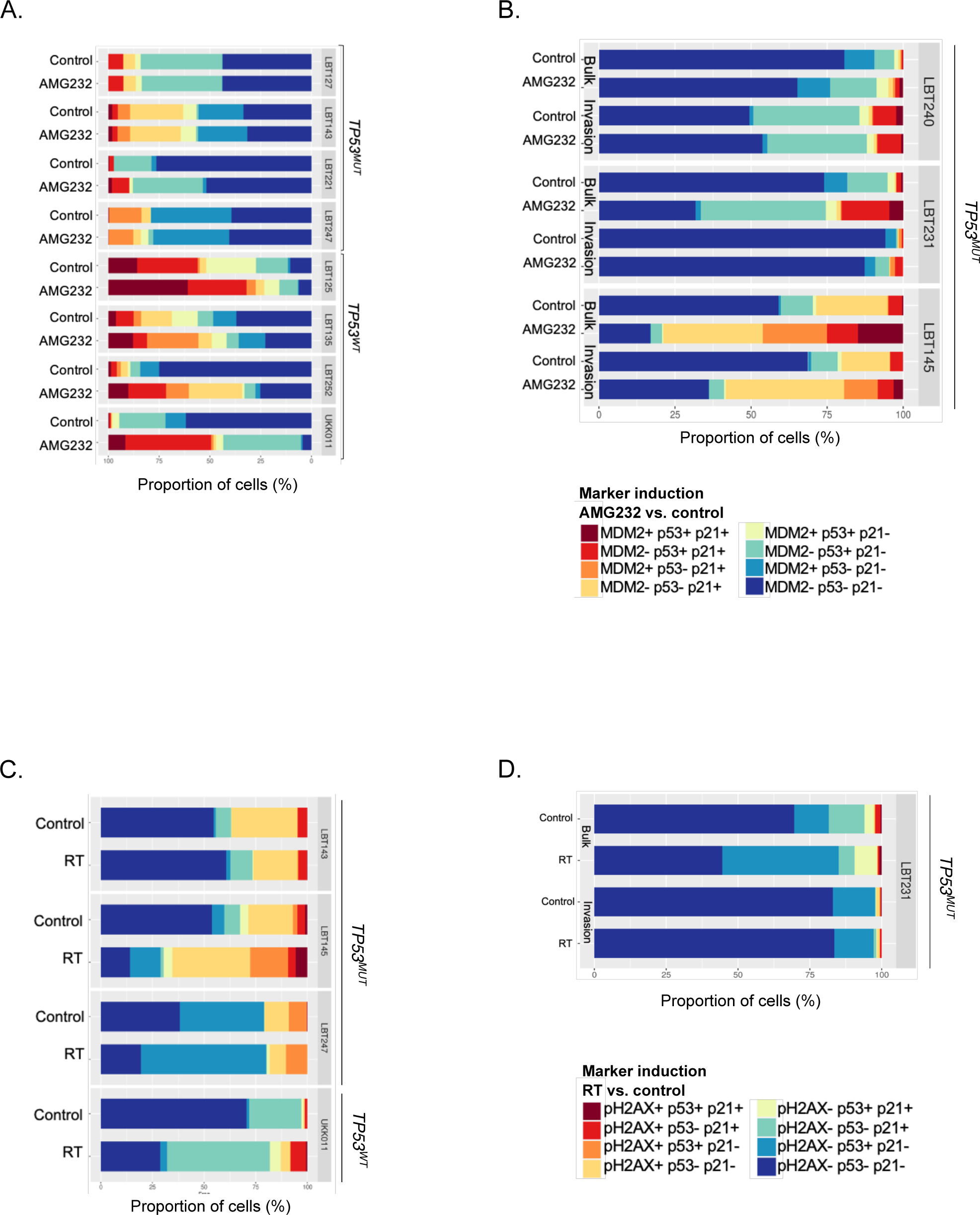

**Supplementary Figure 15.**
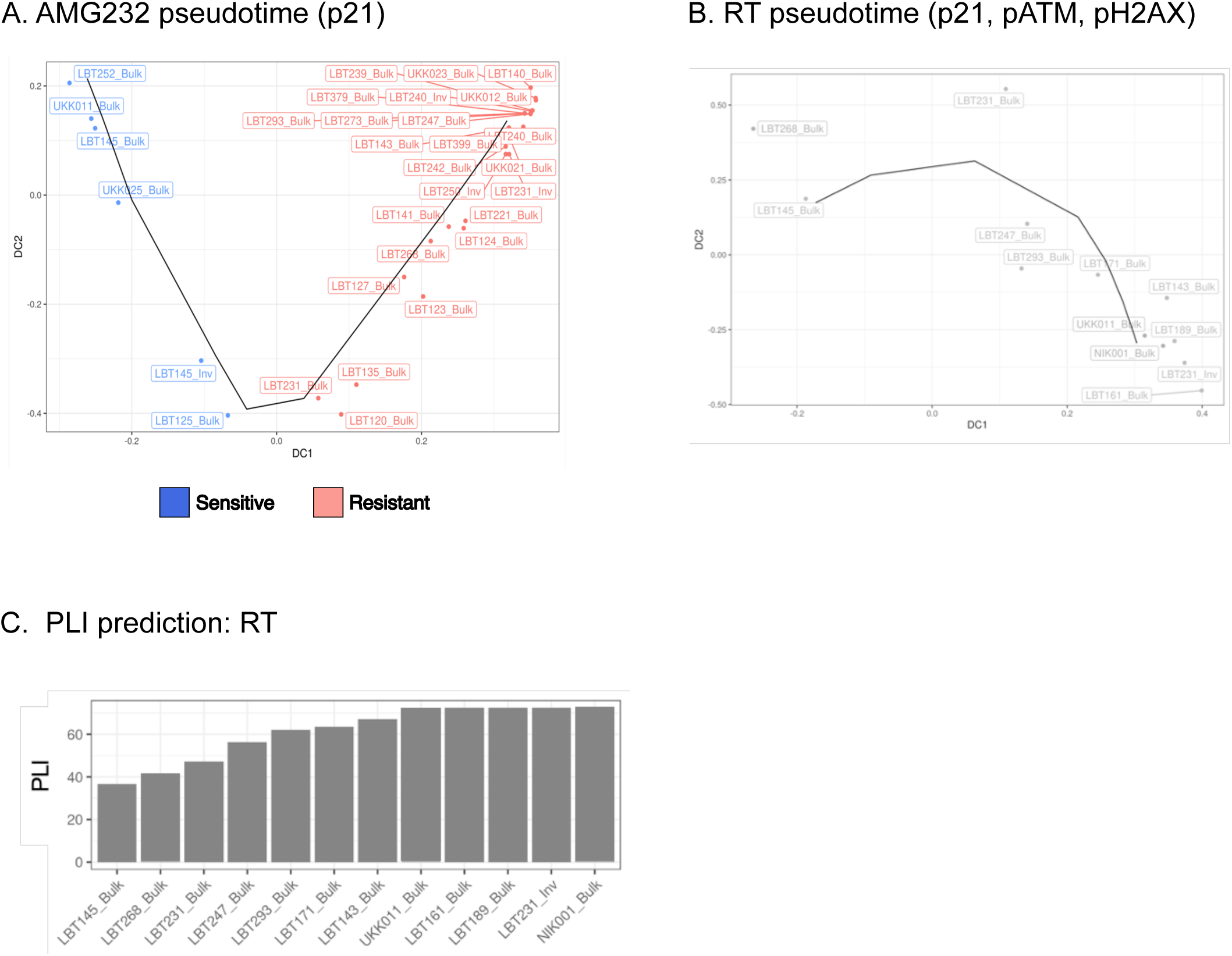

**Supplementary Figure 16.**
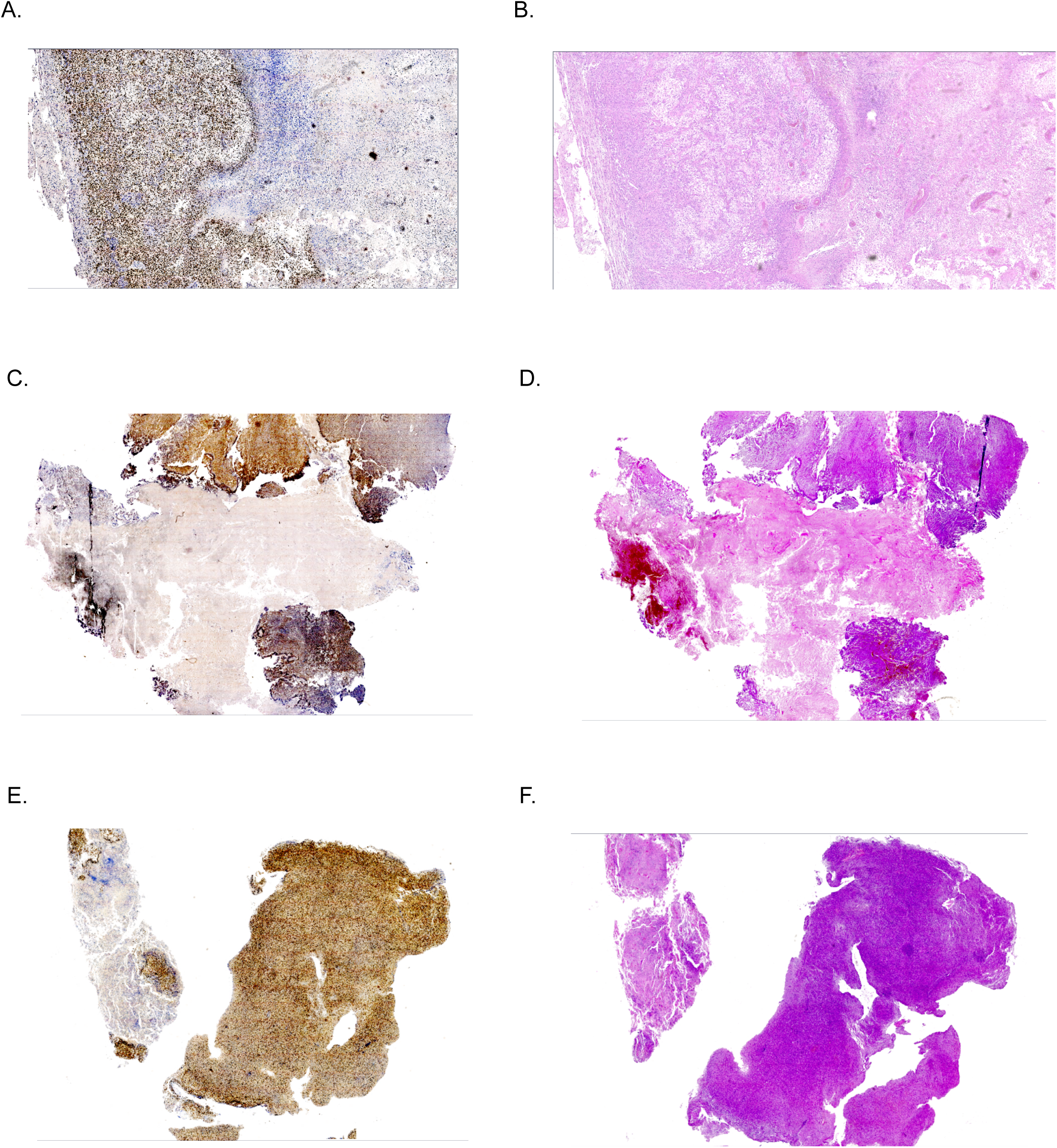

**Supplementary Figure 17.**
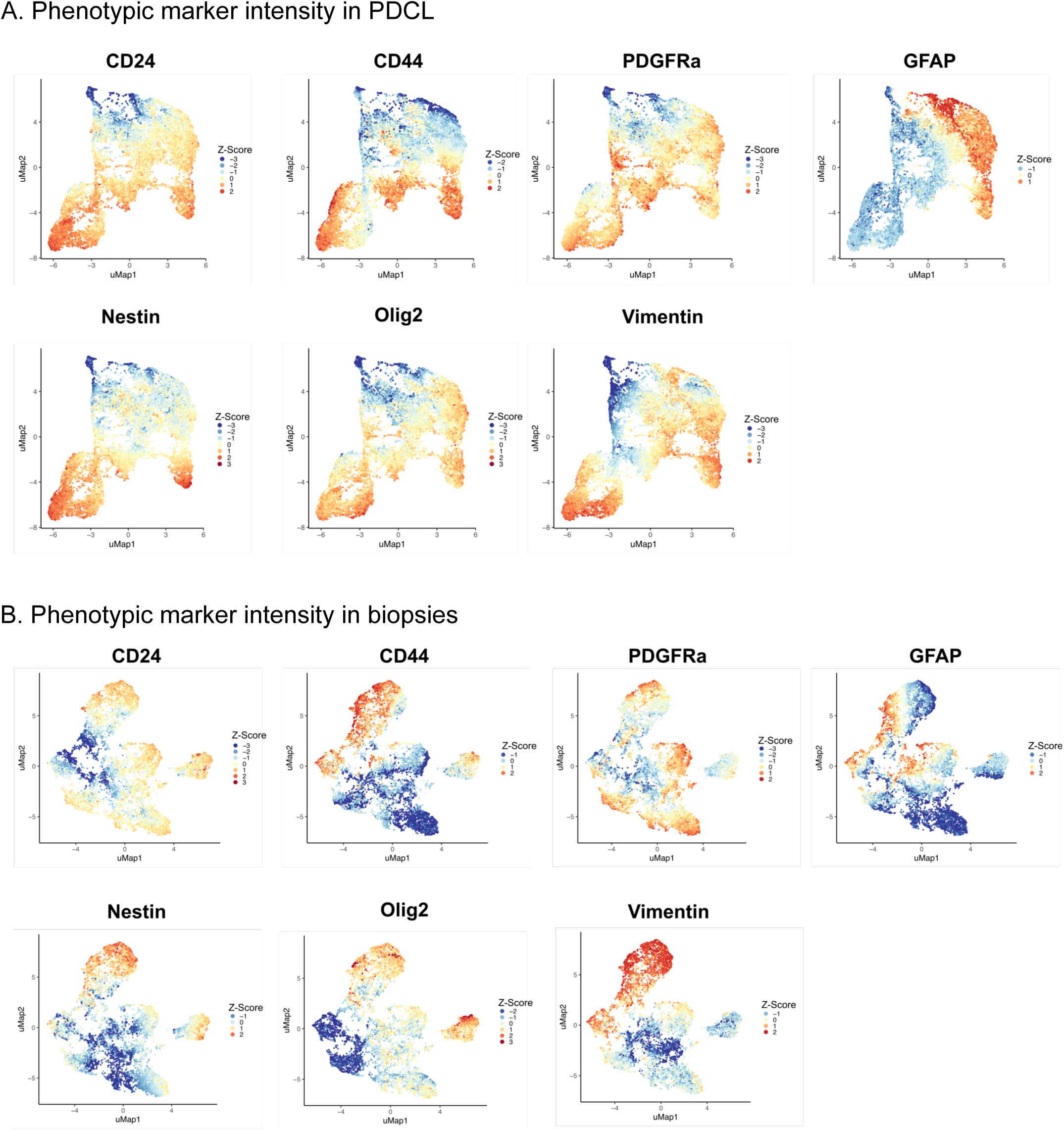

**Supplementary Figure 18.**
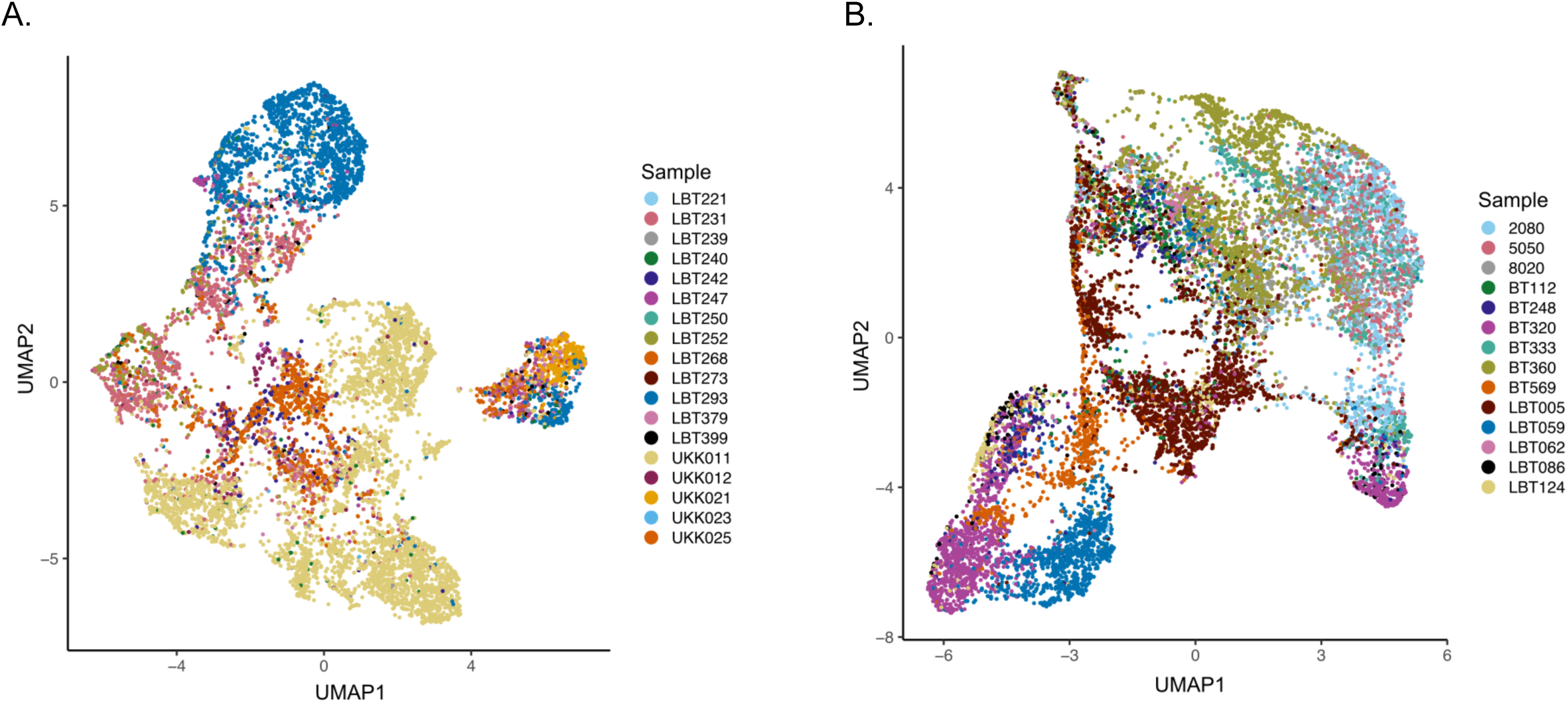

**Supplementary Figure 19.**
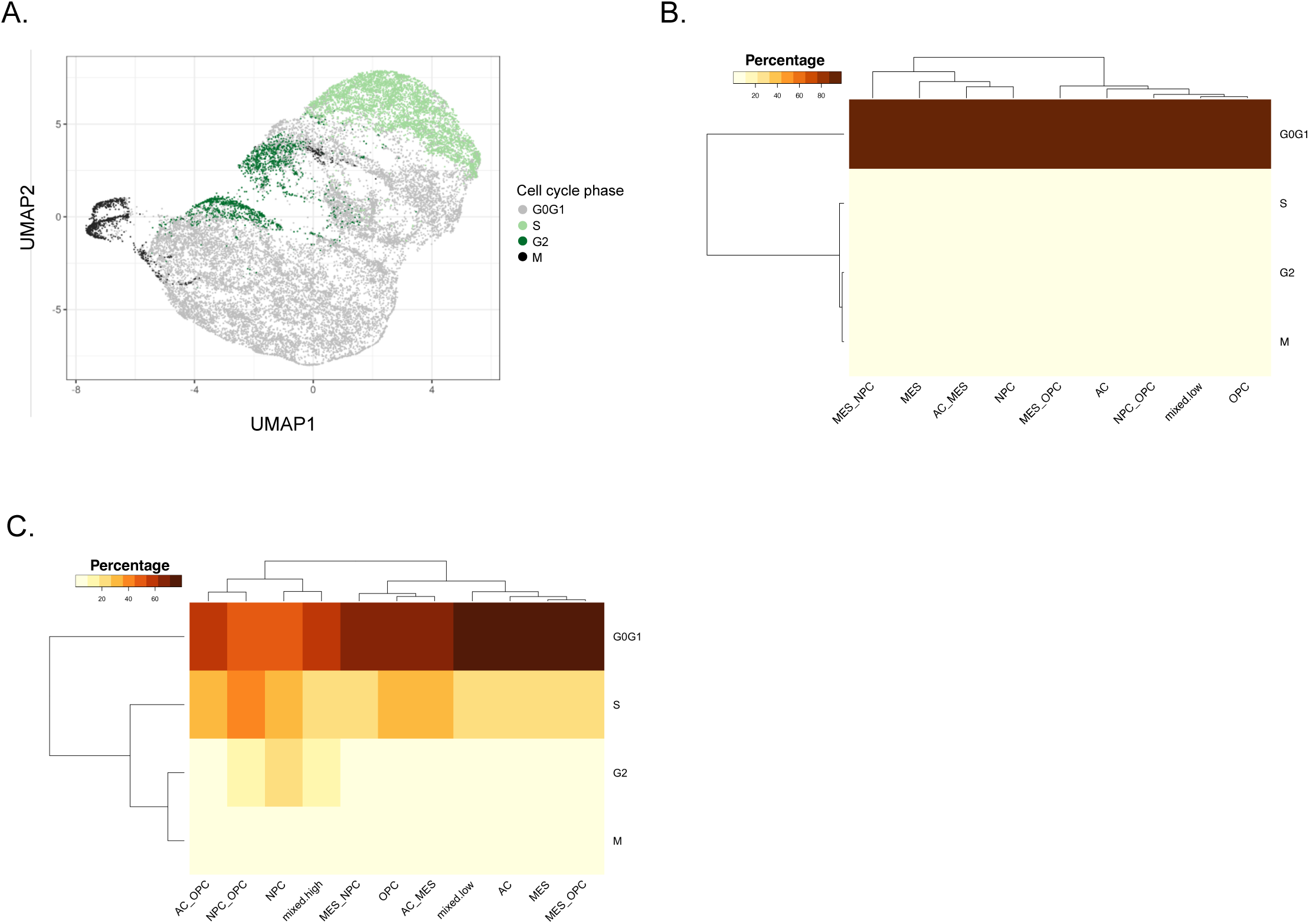

**Supplementary Figure 20.**
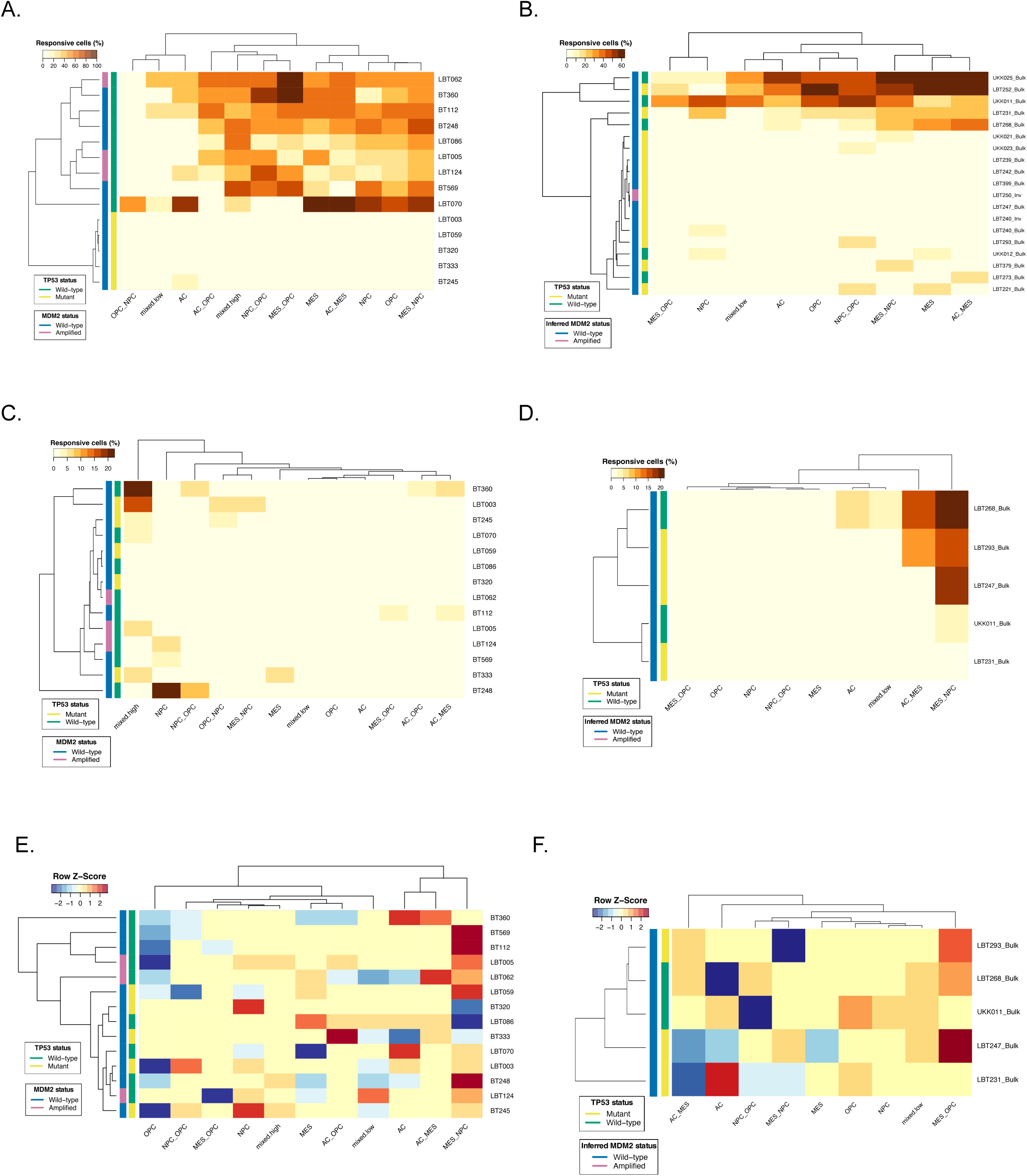

