## Supplementary materials and methods for "Single-cell molecular profiling using ex vivo functional readouts fuels precision oncology in glioblastoma"

### Supplementary Materials and Methods

##### Primary GBM cell dissociation and generation of patiend-derived GBM cell lines

Bulk and invasion specimens of fresh tumor biopsy samples were first weighed separately. Each specimen was then cut with an Aesculap® Disposable Safety Scalpel (VWR) into smaller fragments. Each surgical specimen was separated into 4 fragments: (i) overnight fixation in 4% (v/v) of Pierce™ Formaldehyde (PFA), Methanol-free (PN: 047392.9M, ThermoFischer Scientific) solution; (ii) A snap frozen tumor sample at -80 °C. The remaining parts (≤ 800 mg) were further dissociated for the generation of a patient-derived cell line (PDCL) and/or for direct ex-vivo drug exposure and diagnostic analysis with CyTOF. The samples were dissociated with the MACS Brain Tumor Dissociation Kit (P) (PN: 130-095-942; Miltenyi Biotec) and gentleMACS™ Dissociator. Red blood cells were removed from cell pellets by applying 1 mL of Red Blood Cell Lysis Buffer (PN: R7757, Merck) for 1 min and quenched by prewarmed culture medium (37 °C). The pellets were further purified from myelin and debris by Percoll® PLUS (PN: GE17-5445-02, Merck) gradient separation. PDCLs were cultured as previously described[1,2]. An overview of the 14 PDCLs can be found in Supplementary Table 1.

##### Drug treatments and cytotoxicity assays using AMG232 and radiotherapy

Following dissociation with Accutase, cells were plated (in triplicate for AMG232 (HY-12296, MedChemExpress) and in quadruplicate for RT (RS-2000 Biological Irradiator (Rad Source)) in Corning® Costar® 96-Well Cell Culture Plates at a density of 10 000 cells/well in a volume of 100 µL complete NSA medium per well. After overnight incubation in a humidified incubator at 37 °C, drug dilution series of AMG232 from a stock solution of 10mM were prepared in dimethyl sulfoxide (DMSO) (Sigma-Aldrich) as indicated in the graphs. Each dilution series was performed in triplicate. Similarly, five Corning® Costar® 96-Well Cell Culture Plates were seeded with the PDCLs at a cell density of 10 000 cells/well in a volume of 100 µL complete NSA medium per well. After overnight incubation, each plate was treated with decreasing doses of radiotherapy (starting from 10Gy as the highest one). The control plate was not irradiated and maintained in the incubator. For the time course experiments, cells were harvested at 8, 16, 24 and 48 hours following the start of the treatment, together with a corresponding control. In the mixed experiments, cells from BT333 and BT360 were mixed at different proportions 80:20, 50:50 and 20:80% prior to initial plating. At harvest, samples were prepared for CyTOF analysis (see below). Drug cytotoxicity was analyzed with the CellTiter-Glo® Luminescent Cell Viability Assay (PN: G7572, Promega) at

72 hours after the start of AMG232 treatment and 6 days after RT exposure, according to the manufacturer's instructions. Luminescence was measured in a SpectraMax iD3 Multi-Mode Microplate Reader (Molecular Devices).

Based on the viability experiments, we extrapolated $IC_{50}$ (50% inhibitory concentration), AUC (area under the curve) and PLI (plateau level of inhibition) values. We used the $IC_{50}$-defined concentration (which was ~2.5 μM). The duration of treatment was optimized in time-course experiments, where we prioritized on the time-point where the expression of drug-related markers is maximally altered, while highly preserving cellular integrity.

#### Mouse experiments

Experiments were conducted as previously described[3]. In brief, for the orthotopic intracranial model, before inoculation, BT112 cells were transduced with a GFP/Luciferasee construct. These cells were implanted ($1 \times 10^6$ cells/5 μL) into the brain of $NMRI^{nu/nu}$ Nude mice (8-weeks-old females; (Janvier Labs)). A stereotaxic injection frame was used to inject cells into the right caudate nucleus-putamen (ML +0.15 mm; AP +0.1 cm; DV −0.25 mm). In parallel, from the same culture that was injected into mice, we treated samples with AMG232 using PROSPERO assay, immediately prepared and cryopreserved these samples (10%DMSO/FBS at -80°C).

Animals were imaged weekly using the IVIS Spectrum (Perkin-Elmer; available at the Mosaic Small animal imaging facility, Campus Gasthuisberg, Leuven, Belgium) 10 minutes after injection of 2 mg luciferin (PN: P1042, Promega). When signal of $1x10^8$ photon/second was recorded, the mice were randomly assigned to two experimental groups. The first one (n=3) underwent direct ex vivo analysis. Whole brains were collected, dissociated and directly treated with AMG232 or DMSO (2,5 μM for 16 hours). The second group (n=5), animals were treated by gavage with a single dose of AMG232 (50 mg/kg, n=4) or DMSO (50 mg/kg, n=1). Mice were sacrificed 16 hours post-gavage. Whole brains were once again collected, dissociated and the tissues were fixed and prepared for CyTOF analysis (Figure 6). Samples were stained with the identical panel used for biopsies (Supplementary table 1).

#### CyTOF experiments

##### Sample preparation, treatment and cell harvest

Once cells from each PDCL reached the desired timepoint, cells were collected into 15 mL conical tubes (Sarstedt) by centrifugation (200g, 5 min). After aspiration of the supernatant, cells were washed in sterile DPBS, resuspended in 1 mL of prewarmed StemPro™ Accutase™ Cell Dissociation Reagent (Thermo Fischer Scientific) incubated at 37°C and triturated. Accutase activity was subsequently quenched by the addition of prewarmed complete NSA medium (37 °C), after which cells were again collected by centrifugation (200g, 5 min), washed in DPBS, counted and plated at $1\text{-}2x10^6$ cells per Corning T25 cell culture flasks. In this project, samples for CyTOF samples were treated with DMSO, AMG232 and RT at the indicated doses; in parallel, the same batch of cells was subjected to cytotoxicity assay (see above). After incubation of the samples for the indicated duration, samples were collected into 15 mL conical tubes (Sarstedt) and centrifuged (200g, 5 min). After aspiration of the supernatant, cells were washed in 5 mL of prewarmed DPBS (37 °C) (Thermo Fischer Scientific) (1200 rpm, 5 min). In order to dissociate cell clusters and ensure single-cell suspensions, pellets underwent once again an Accutase treatment with StemPro™ Accutase™ Cell Dissociation Reagent (Thermo Fischer Scientific).

**IdU staining**

Subsequently, the single cell suspensions were placed in 1ml pre-warmed medium and treated with IdU (Cell-ID™ 127 IdU, PN: SKU201127, Fliudigm) at a final concentration of 50μM for 30 minutes at 37°C, to stain for S-phase cells. This procedure was quenched by adding DPBS (5-times volume of the cell suspension) and washed twice with DPBS at 200g for 5 minutes.

**Cisplatin viability staining**

In order to discriminate dead cells from live cells, samples were stained with Cell-ID™ Cisplatin (PN: SKU201064, Fluidigm) at a final concentration of 0,1 μM. The samples were mixed by vortexing and incubated for 3 min at RT. Cisplatin staining was quenched by the addition of 5 mL of Maxpar® Cell Staining Buffer (PN: SKU201068, Fluidigm) after which the samples were centrifuged (200g, 5 min) and the supernatant was discarded.

**Fixation and cryopreservation**

Immediately after Cisplatin staining the cells were fixed by resuspending them in 1 mL of Pierce™ Formaldehyde (PFA), Methanol-free (Thermo Fischer Scientific) solution at a final concentration of 2% (v/v) in DPBS. The samples were incubated for 30 min on a spinning wheel at room temperature. From this point, all centrifugation steps were performed at higher rotation speeds (800 x g, 5 min) to optimize cell recovery. After fixation, samples were washed twice with Maxpar Cell Staining Buffer. After aspiration of the supernatant, cells were resuspended in Heat-Inactivated Fetal Bovine Serum (FBS) (Gibco) and cryopreserved at -80 °C in a 2 mL round bottom cryovials (VWR) supplemented with 10% of DMSO (Sigma-Aldrich). The following day, CyTOF-ready samples were transferred into liquid nitrogen tanks[4].

**Barcoding**

Thawed PDCLs, tumor samples and PDX probes were immediately transferred into cold washing solution (equal volumes of 1% BSA, Cell Staining Buffer (Fluidigm) to which we added Benzonaze (25U/ml Ultra-pure, >250 units/μl, >99%, from a 25 KU stock (PN: E1014-24KU, Sigma-Aldrich)). The samples were washed twice with DPBS prior to barcoding. As multiple samples were intended to be processed simultaneously, each sample was barcoded individually and pooled together in order to reduce the total reaction volume, minimize sample loss due to multiple washing steps and technical variability between the samples. This way, subsequent antibody staining procedures could be performed on multiplexed sample mixtures, which are maximally composed of 20 uniquely barcoded probes. Here, the Cell-ID™ 20-Plex Pd Barcoding Kit (PN: 201060, Fluidigm) was applied according to the manufacturer's instructions through which each sample received a unique barcode. Following barcoding, samples were resuspended in 100 µL of Maxpar® CSB (Fluidigm), and combined into a single Corning™ Falcon™ Test Tube with Cell Strainer Snap Cap (Thermo Fischer Scientific) after which the multiplexed sample was centrifuged (800 x g, 5 min) and the supernatant was discarded.

**Antibody cocktail preparation**

The panel mainly contains antibodies ordered by Fluidigm. Antibodies against markers which were not available at Fluidigm, were obtained by different companies, custom-labelled and tested/optimized at our lab by using appropriate controls (see Supplementary Table 2). For each tube, the appropriate antibody cocktails were prepared, i.e. one to stain cell surface and intracellular (cytoplasmic and nuclear) markers and one to stain phosphorylated protein markers (see supplementary Table 2). For each antibody an adapted dilution was added to a total volume of 50 µL antibody cocktail per individual sample. Cell surface and intracellular antibody cocktails were prepared in DPBS (Thermo Fischer Scientific) supplemented with 0,2% (v/v) Triton™ X-100 (Sigma-Aldrich) and 1% BSA (w/v) Fraction V (Biotium), whereas phosphoprotein antibody cocktails were prepared in Maxpar Cell Staining Buffer (Fluidigm). Each antibody cocktail was filtered through Centrifugal filters (0,1 µm Ultrafree 500 µL-MC, PN: UFC30VV25, Milipore) at 12,000 x g for 4 minutes. Staining of intracellular/cell surface and phosphoprotein markers was carried out separately and consecutively.

**Cell surface and Intracellular staining**

Prior to antibody staining, the multiplexed samples (PDCLs and tumor samples) were washed and incubated for 30 minutes at room temperature in DPBS containing Triton X-100 and 1% BSA. After the Triton X-100 permeabilization, the tumor samples were blocked by Heparin sodium salt (PN: H3393-25KU, Sigma-Aldrich) for 20 min at room temperature which was followed by an additional blocking step with 20µL FcR blocker (PN: 130-059-901, Miltenyi Biotec, 10 minutes at 4°C); the PDCL samples directly underwent cell surface/intracellular staining. 50 µL of antibody cocktail per individual sample was added to each test tube and incubated for 30 minutes at room temperature, while the tubes were occasionally inverted, after which the cell pellets were washed with Maxpar® CSB (Fluidigm).

After the second wash, the supernatant was removed and the pellets were placed on ice for 15 minutes anticipating the next staining round.

**Phosphoprotein staining**

Phosphoprotein staining was initiated with the resuspension of the cells in 1 mL of 4°C 90% (v/v) Methanol GPR Rectapur® (VWR) followed by a 5 min incubation on ice. After centrifugation of the sample (800 x g, 5 min) and aspiration of the supernatant, the cells were washed twice in 2 mL of Maxpar® CSB (Fluidigm). After aspiration of the supernatant, the phosphoprotein antibody cocktail was added to the multiplexed sample, with 50 µL of antibody cocktail per individual sample. After incubation for 30 minutes at room temperature with regular mixing, the sample was centrifuged (800 x g, 5 min) and the supernatant was discarded. Next, the sample was washed in 1 mL of Maxpar® CSB (Fluidigm) (800 x g, 5 min) to remove any unbound antibody. After aspiration of the supernatant, the cells were resuspended in 1 mL of intercalation solution for each sample by adding Cell-ID™ Intercalator-Ir (PN: 201192A, Fluidigm) into Maxpar® Fix and Perm (PN: 201067, Fluidigm) to a final concentration of 125 nM (1,000X dilution of the 125 µM stock solution) and vortex to mix. The pellets were resuspended and stored overnight at 4 °C. Prior to sample acquisition on the Helios™ Mass Cytometer (Fluidigm), the samples were washed twice with Maxpar® CSB (Fluidigm) (800 x g, 5 min) supplemented with Benzonase (25U/ml) and once with Maxpar® Cell Acquisition Solution (CAS, PN: 201237, Fluidigm). Once the pellets were ready for acquisition, the cells were resuspended in (1x) EQ Four Element Calibration Beads (PN: SKU 201078, Fluidigm) diluted in CAS reaching $3 \times 10^5$ cells/ml.

**CyTOF data acquisition**

The data was acquired on a Helios® Mass Cytometer (Fluidigm) in FCS format (the KU Leuven Flow and Mass Cytometry Facility at Campus Gasthuisberg, Leuven (Belgium)). CyTOF software version 6.7.1016 and separate PDCLs, patients' and PDX sample templates were used to acquire and normalize data from the stained samples. The PDCL sample pool was separated and analyzed in three different batches. Data of batch 1 was collected on a different date than batch 2 and 3 and therefore the phenotypic analysis of these samples was performed separately from batch 2 and 3 (further details below). From the biopsy samples we have two datasets available, whereby only the phenotypes were mapped only in the latest dataset, while in the old one we only evaluated drug responses of SOX2+ tumor cells. For each one, batch-normalization samples were included (BT112 and BT245 - control and AMG232-treated samples) Cell count and batch annotation for each sample are included in Supplementary Table 4.

**High-dimensional data preprocessing**

**FCS debarcoding**

Normalized data from each tube was collected and written in a barcoded FCS which needed to be deconvoluted into separated FCS files of each barcoded population. The tool used for debarcoding the palladium stained samples was the Matlab Single Cell Debarcoder v0.2, which besides the single-cell deconvolution algorithm applies an additional doublet filtering scheme[5].

**Quality control**

The raw data files were already normalized for technological and batch effects by the use of EQ normalization beads and batch-correction samples (BC samples: BT112 and BT245 - control and AMG232-treated), which were run on the Helios™ Mass Cytometer (Fluidigm) in parallel within each multiplexed tube of PDCL or patient samples. Next, the FCS files were loaded into the online tool Cytobank for QC. Data quality was confirmed through visual inspection of the course of the bead signals over time. This way, a homogeneous course of the bead signals over time was considered indicative for experiments that ran without major disturbances in time such as clogging, speed changes or air pressure measurements when the tube is empty. Additionally, event length, representing the time for single cells (singlets), doublets or multiples to reach the detector, was visually inspected as well as part of QC.

**Automated gating**

After QC, the data was automatically gated in Maxpar® Pathsetter™ (version 2.0.45) through a tailored model that was developed and trained for the purposes of this project, i.e. GBM cells. The gating strategy applied eight consecutive layers of stringency. This way, live single cells were selected through the application of a tailored gating window. Starting from the ungated population, cells were sorted out from beads by plotting the bead signals over time. In the next gating rounds, single cells or singlets were selected against doublets by inspecting the following parameters over time, in a consecutive manner: residual, offset, center, width and event length. In the final and crucial gating step, the viable portion of the single cells was selected as the subpopulation with high DNA1/DNA2 content (193Ir) and low cisplatin levels. From here Maxpar® Pathsetter™ generates cleaned FCS files and a report summarizing QC metrics for acquisition and modeling quality for each population, cell number, percent of total cells extracted from each parent population.

#### High-dimensional data analysis

**Batch normalization**

Intensity values were asinh transformed. Four samples were measured in all batches as technical replicates to correct for technical sources of variability. To that end, we estimated the transformation function following equation eq.1 that maximized the overlapping coefficient between the density functions of the individual samples belonging to different batches.

Eq1 → y' = alpha + beta*y

This approach was repeated independently for every marker in every batch. Batch 3 was selected as reference since it contained the largest number of cells. Asinh values were further transformed to z-scores (Supplementary Figure 1C and D).

**Population identification and cellular abundance in patient samples**

Two separate datasets, acquired at different times with different batches of antibodies, were included in the analysis. The analysis pipeline begun with the isolation of the tumor cells (SOX2+) from the immune cells (CD45+/CD3+ and CD68+) by manual gating, a visual inspection of a series of two-dimensional scatter plots (Supplementary Figure 12).

**Tumor subtype annotation and identification of phenotypic patterns in PDCLs and biopsies**

Clustering was performed exclusively in a subset of 10,000 cells that was selected after stratified proportional sampling. Identified clusters were mapped to known cell phenotypes following manual annotation. This way, for each cell we have 4 annotations. If the annotation of 2 or more clustering methods agreed, then the cell was labelled with the most common annotation. If two annotations had the same number of hits, the annotation given by Clara[6] was prioritized. If all four clustering methods disagreed on the given annotation, the cell was labelled as "noise". To predict the rest of the cells not included in the clustering, we trained a support vector machine (SVM) with a radial kernel following a 3-times repeated 10-fold cross-validation scheme.

The marker expression values in the patients' samples were asinh transformed and then normalized by z-scoring, after which phenotypic clustering of the tumor subset was performed only by PhenoGraph[7], on a subset of 25.000 cells of the new biopsy datasets, based on the expression of seven tumor markers (Vimentin, Nestin, CD44, CD24, Olig2, PDGFRa, GFAP). This analysis generated heatmaps of 18 cluster which were all annotated to a specific subtype (mesenchymal, astrocytic, oligodendrocytic, neural) and transitional states (astrocytic /mesenchymal, mesenchymal /neural, mesenchymal / oligodendrocytic, neural / oligodendrocytic). Undifferentiated tumor progenitors not expressing clearly any of the above-mentioned markers were assigned as "mixed.low". After the annotations were mapped to each cluster, a Support Vector Machine (SVM) with the same settings was trained and applied on one of the whole dataset. In this way, the whole dataset was annotated. This allowed us to observe the distribution of the clusters not only in baseline, but also in the different therapy conditions (AMG232 and RT) (Figure 5, Supplementary Figure 17, Supplementary Figure 18, Supplementary Figure 20).

**Unbiased probability modeling of drug responses in PDCLs using AUC and PLI**

The induction of functional markers p53, p21, MDM2, pH2AX, BAX, CC3, and pATM was evaluated for the different treatments. Here, a cell was defined as induced if its expression for a given marker was significantly higher than the distribution defined by the untreated cells (background distribution). These background distributions were modelled as normal distributions for each marker/sample. The cutoff for induction was defined in the 95$^{th}$ quantile of the distribution (p-value < 0.05). In the case of p53, since p53-mutant cell lines showed bimodal distributions in the untreated samples, these were previously deconvolved using Gaussian Mixture Models (GMMs). After deconvolution, the distribution with the lowest average was used to model the background distribution (Supplementary Figure 3; Supplementary Figure 13). The total induction of each marker in each sample was then estimated as follows:

$$total\_induction_{i,j} = treatment\_induction_{i,j} - control\_induction_{i,j}$$

where treatment_induction is the percentage (%) of cells induced for sample i and marker j upon treatment, and control_induction is the % of cells induced for sample i and marker j in the untreated sample (Figure 2G and I, Supplementary Figure 2E and F, Supplementary Figure 14).

These fingerprints were then used to optimize a predictive panel of treatment response. For each treatment, a subpanel of functional markers was selected based on prior knowledge: p21, p53, MDM2, BAX, and CC3 for AMG; p21, p53, MDM2, BAX, CC3, pATM, and pH2AX for RT. An exhaustive optimization scheme was followed where panels of size 1, 2, …, N (N=total number of markers) were trained. For each panel size, all the possible marker combinations were evaluated. For each marker combination, a fingerprint for each sample was built with the percentage of cells showing a specific induction profile. For example, when evaluating a panel based on p21 and p53 the fingerprint has four features (p21 low p53 low, p21 high p53 low, p21 low p53 high, and p21 high p53 high). Here, high stands for the cells that showed induction whereas low stands for the cells that did not show an induction (Supplementary Figure 4). These induction profiles were dimensionally reduced using diffusion maps (DM) as implemented in the R package Desntiny[8] (Supplementary Figure 5, Supplementary Figure 9A and G, Supplementary Figure 15A and B). A trajectory was then defined in the reduced space using Slingshot[9]. The pseudotime derived from trajectory analysis was correlated with therapy response using two different endpoints (AUC and PLI) (Figure 3; Supplementary Figure 6; Supplementary Figure 9B-C, E-F, H-I and K-L; Supplementary Figure 10D-H; Supplementary Figure 11B-F). The correlation scores were further used to build a Pareto front (Figure 3). Briefly, for each panel size, we selected the panel combination giving the best result. Then we compared the panel performance (correlation scores, y-axis) as a function of panel complexity (number of markers, x-axis). Following Occam´s razor principle, we selected the simplest model that explains the data by applying the elbow criterion to the Pareto front (Figure 3).

##### Analysis of the cell cycle phases in PDCLs and patients' samples

The markers used for defining cell cycle phases were: pH3 (mitosis, M-phase), IdU (synthetic, S-phase), CNNB1 (G2-phase), pRB1 (G0/G1) and Ki67 (proliferation)[10] and the population identification was performed by manual gating (Supplementary Figure 19).

##### Projection of the predictive model on patients' samples

The total induction of all the markers were used to build a functional fingerprint for each sample in each treatment. These functional profiles were used to predict drug-response related end-points, i.e., area under the drug-response curve (AUC), and maximum inhibition. We exhaustively trained unsupervised models using subpanels of the functional profiles of size 1:n, being n the number of functional markers considered (5 in the case of AMG and 7 in the case of RT). For each panel, we estimated a trajectory using pseudotime analysis (slingshot) in a dimensionally reduced space that was calculated using diffusion maps. This trajectory was then linearly correlated to the drug-response endpoints and, for each model, the corresponding Pearson correlation (R) and p-value were extracted.

For each panel size, the marker/combination of markers yielding the best performance was used to build a Pareto-front with the x-axis representing the panel complexity (number of markers) and the y-axis representing the panel performance (p-value/R). The optimal panel was selected applying the elbow criterion. The optimal panels for each treatment were further explored to build more accurate models of drug-response prediction. These fine-tuned models were used to project the biopsy samples and give an estimation on their drug-response. The same approach was used using only the control samples.

##### Ex vivo PDX data analysis

Similarly, as in the biopsy dataset, tumor cells (SOX2+) were isolated by manual gating when plotted against CD45+ cells. The functional analysis was performed by GMM modeling and adjusting the expression cut-offs, as described above.

#### Code availability

All codes are provided on the github page, which will be made available upon publication of the manuscript.

##### Figure preparation

The graphical abstract and schematic diagrams were prepared in BioRender.com. Statistical analysis, generation of dose-response viability curves and analysis was performed with GraphPad Prism 9.4.1 (458) Macintosh Version by Software MacKiev © 1994-2022 GraphPad Software, LLC.

**Supplementary Figure Legends:**

**Supplementary Figure 1** GBM pathophysiology assessed in 14 patient-derived cell lines. (a) Most common genetic abnormalities measured by whole-genome sequencing (WGS). (b) qPCR expression signatures. (c) Heatmap overview of protein landscape of the phenotypic markers as measured by CyTOF. (d) Density plots of the baseline expression of phenotypic markers measured through CyTOF. (e) Correlation matrix between RNA expression (bulk RNA sequencing) vs protein expression (CyTOF).

**Supplementary Figure 2** Overall responses of PDCL pool, viability and single-cell molecular profiles of BT360/BT333 sample mixtures. (a) Bar plot of AUC values representing the cell viability after AMG232 treatment (0-10 µM for 72 hours). (b) Bar plot of AUC values representing the cell viability after radiotherapy exposure (0-10 Gy for 6 days). The values represent mean ± SD of three (AMG232) or four (RT) replicates and were normalized to 100% assigned to the vehicle control for each assay. Overall responses of the models are represented as AUC values and stratified by *TP53* mutational status (*TP53*$^{MUT}$- red dots; *TP53*$^{WT}$ blue dots and *MDM2*$^{AMP}$ – blue stripes). Wilcoxon rank sum test was used to calculate statistical significance between the *TP53*$^{WT}$ and *TP53*$^{MUT}$ groups (ns = not significant ($P > 0.05$); *$P \leq 0.05$; **$P \leq 0.01$; ***$P \leq 0.001$; ****$P \leq 0.0001$). (c) Viability treatment with AMG232. (d) Dose response curves after irradiation of cell mixtures of BT360 and BT333 at 80-20%, 50-50% and 20-80%. (e) Stacked bar plot representing drug-induced signatures (MDM2-p53-p21) upon AMG232 treatment. (F) Stacked bar plot representation of drug-induced signatures (pH2AX-p53-p21).

**Supplementary Figure 3** Normalized z-scored density plots representing expression changes upon treatment of PDCLs and mixed samples. (a) Drug-related markers are shown and the definition of expression cut-offs is explained in Supplementary Methods section. (b) Normalized z-scored density plots of p16- and PDGFRa-expression in the PDCL pool.

**Supplementary Figure 4** Panel optimization for unbiased identification of most optimal therapy-dependent marker combination based on AUC/PLI extrapolated from cytotoxicity treatments upon AMG232 (readout at 2.5 µM) and RT (10 Gy). Literature-guided signatures are highlighted in blue bars and optimized (selected) signatures in green bars.

Upon AMG232 treatment: (a) Marker combination correlated (R) to AUC. (b) Marker combination correlated (R) to PLI.

For RT: (c) Marker combination correlated (R) to AUC. (D) Marker combination correlated (R) to PLI.

**Supplementary Figure 5** Dimensionality reduction of optimized response scores of each cell line into diffusion maps. (a) Map of p21 response score of each cell line upon AMG232 treatment. (b) Diffusion map of cell lines upon AMG232 treatment, whereby *MDM2*$^{AMP}$ cell lines are highlighted. (c) Diffusion map generated by the response scores (pH2AX-pATM-p21) of the PDCLs upon RT. (a) and (c) Based on long-term viability response, the PDCLs are colored by distinct color code indicating sensitive, intermediate (mixed) or resistant.

**Supplementary Figure 6** Trajectory inference (pseudotime) of response scores and cell survival metrics (AUC/PLI) upon AMG232 and RT. Based on long-term viability response, PDCLs are colored by distinct color code indicating sensitive (blue), intermediate / mixed (green) or resistant (red).

Upon AMG232: (a) Pseudotime axis of p21-response scores plotted against AUC values for each cell line. (b) Highlighting *MDM2*$^{AMP}$ cell lines which skew the linearity of the model. (c) Pseudotime of p21 response scores plotted against PLI values for each cell line. (d) Improvement of the linearity/smoothness of the continuous path after stratification of *MDM2*$^{AMP}$ cell lines. Upon RT: (e) Pseudotime trajectory of (H2AX-pATM-p21) response scores plotted against AUC values for each cell line. (d) After training the predictive model, AUC values were plotted against Predicted AUC.

**Supplementary Figure 7** Cell cycle annotation of two consecutive PDCL batches. (a) UMAP (of 11/14 PDCLs analyzed in batch 2 and 3) representing annotated cells in each cell cycle groups. (b) Expression levels of markers used for manual annotation of each cell cycle phase. (c) UMAP representation of the cell cycle phases (of 3/14 PDCLs analyzed in batch 1). (d) Expression levels of markers used for manual annotation of each cell cycle phase.

**Supplementary Figure 8** Treatment effects on the distribution of the cell cycle phases in PDCLs. (a) Boxplots overlaid with dot plots, where each dot represents *TP53*$^{MUT}$ cell line. The percentages of tumor cells from each *TP53*$^{MUT}$ cell line are represented in each cell cycle phase and treatment condition. (b) Boxplot overlaid with dot plot, where each dot is a *TP53*$^{WT}$ cell line. The percentages of tumor cells from each *TP53*$^{WT}$ cell line are plotted in each cell cycle phase and treatment condition. The significance of the difference between both treatments vs the control was determined by Wilcoxon Rank-Sum test: *p≤0.05, **p≤0.01, ***p≤000.1. (c) Stacked barplots representing on the distribution of cell cycle phases in p21-induced cell populations (responsive population in each PDCL) vs p21-repressed (unresponsive population in each PDCL) in AMG232-treated samples. (d) Stacked barplots representing on the distribution of cell cycle phases in irradiated PDCL samples. The top four signatures are considered as responsive to RT, in contrast to the four profiles below.

**Supplementary Figure 9** Pseudotime model predicting AUC/PLI values based on cell cycle signatures upon AMG232 (a-f) and RT exposure (g-l). The color code indicates sensitive (blue), intermediate/mixed (green) and resistant (red) responses recorded after long-term exposure in AMG232 viability assay. (a) Diffusion map of PDCL-axis ordered by cell cycle scores (control vs AMG232). (b) Linear correlation of AUC and pseudotime. (c) Correlation between AUC and predicted AUC for the cell cycle scores. (d) Box plots overlaid with dot plots, where each dot represents the pseudotime value of each PDCL. The pseudotime values are compared between each PDCL-response class (sensitive, mixed, resistant). (e) Correlation between pseudotime value of each PDCL and PLI. (f) Correlation between PLI and Predicted PLI. (g) Diffusion map of PDCL-axis ordered by cell cycle scores (control vs RT). The color code indicates sensitive (blue), intermediate/mixed (red) and resistant (green) responses recorded after long-term exposure in a RT viability assay (h) Linear correlation of AUC and pseudotime. (i) Correlation between AUC and predicted AUC for the cell cycle scores. (j) Box plots overlaid with dot plots, where each dot represents the pseudotime value of each PDCL. The pseudotime values are compared between each PDCL-response class (sensitive-green, mixed-blue, resistant-red). (k) Correlation between pseudotime value of each PDCL and PLI. (l) Correlation between PLI and Predicted PLI. The significance of the difference between pseudotime values for both treatments vs the response class was determined by Wilcoxon Rank-Sum test: *$p \leq 0.05$, **$p \leq 0.01$, ***$p \leq 000.1$.

**Supplementary Figure 10** Pseudotime analysis of control samples focused on prediction of survival after AMG232 treatment. (a) Panel optimization to identify the most predictive marker of PLI (R). (b) Panel optimization to identify the most predictive marker of AUC (R). (c) Paretofront of number of markers vs R-value correlating to AUC (full line) / PLI (dashed line). Here, p53 was selected the following analysis. (d) Correlation of baseline p53 expression in control samples with AUC, generating a pseudotime value for each PDCL. (e) Correlation between pseudotime score and AUC. (f) Correlation of baseline p53 expression in control samples with PLI, generating a pseudotime value for each PDCL. (g) Correlation between pseudotime score and PLI. (h) Correlation between PLI and predicted PLI. The color code indicates sensitive (blue), intermediate/mixed (red) and resistant (green) responses recorded after long-term exposure in AMG232 viability assay. (i) Boxplot showing difference in p53 expression between $TP53^{MUT}$ and $TP53^{WT}$ PDCLs. (j) Boxplot showing difference in p53 expression between $TP53^{MUT}$ and $TP53^{WT}$ freshly isolated patient’s samples.

**Supplementary Figure 11** Pseudotime analysis of PDCL-control samples focused on prediction of survival after RT treatment. (a) R-correlation vs marker combination for both AUC (full line) and PLI (dashed line). Here, pATM was selected for the downstream analysis. (b) Pseudotime values generated based on correlation between pATM expression and AUC. (c) Correlation between the pseudotime score and AUC. (d) Correlation between PLI and predicted PLI in baseline samples upon RT. (e) Correlation between PLI and pATM expression. (f) Correlation between PLI and pseudotime (pATM). The color legend indicates responsiveness classes determined by the survival profile of each cell line upon long-term RT exposure in viability assays. Sensitive PDCLs (blue), mixed / intermediate (red) and resistant (green).

**Supplementary Figure 12** Cell lineage tracing and assessment of the distribution of three major cellular components in GBM tumor resections: malignant cells (SOX2+), T cells (CD3+) and myeloid cells (CD68+). (a) Subsampled UMAP embedding of the three cellular compartments. (b) UMAPs colored by the normalized, z-scored expression levels of phenotypic markers used for identification of each population. (c) Relative distribution of each cell population in biopsy samples.

**Supplementary Figure 13** Normalized, z-scored density plots representing expression changes upon treatment of biopsy samples. (a) Drug-related markers are shown and the definition of expression cut-offs is explained in Supplementary Methods section. (b) Normalized, z-scored density plots of p16- and PDGFRa-expression in the biopsy pool.

**Supplementary Figure 14** Expression patterns of literature-guided drug-related markers among distinct biopsy samples in control and treatment conditions represented as stacked barplots. (a) Differences in responses to AMG232 perturbation between $\textit{TP53}^{WT}$ and $\textit{TP53}^{MUT}$ samples. (b) Stacked barplots showing the percentage of cells with specific marker expression among three patients' biopsies from which bulk and invasion samples were treated with AMG232. (c) Differential response signature expression in RT-treated patient biopsies. (d) Protein expression patterns in control and RT treated samples (tumor core and invasion front) derived from a single patient.

**Supplementary Figure 15** Response prediction (PLI) of patients' biopsies. (a) Diffusion map of the response score (p21) of patients' samples generating a pseudotime path between sensitive (blue) and resistant (red) samples. (b) Diffusion map of the response score (pH2AX-pATM-p21) of patients' samples generating a pseudotime line. (c) Barplot of predicted PLI for RT-treated biopsies.

**Supplementary Figure 16** Representative IHC images showing heterogeneous p53 expression and corresponding H&E staining in few patient-derived brain tumor tissues. Scale bars were set at 20μm magnification. (a, b) p53 expression and H&E staining in patient LBT120. (c, d) p53 expression and H&E staining in patient LBT145. (e, f) p53 expression and H&E staining in patient LBT252.

**Supplementary Figure 17** Inter- and intra-tumor heterogeneity in GBM sample pools. (a) UMAP representation of expression of phenotypic markers (OLIG2, PDGFRa, CD44, Nestin, Vimentin, CD24, GFAP) used to annotate tumor subpopulations in PDCLs. (b) UMAP representation of expression of phenotypic markers used to annotate tumor subpopulations in patients' biopsies.

**Supplementary Figure 18** UMAP visualization of control (vehicle treated) sample pools. (a) Colored by patients' samples. (b) Colored by PDCLs.

**Supplementary Figure 19** Cell cycle annotation in patients' samples and evaluation of the distribution of cell cycle phases across tumor cell subpopulations. (a) UMAP embedding of patient-derived tumor cells colored according to the cell cycle phase. (b) Heatmap showing the percentage distribution of each cell cycle phase across phenotypic clusters in patients' tumor samples. (c) Heatmap showing the percentage distribution of each cell cycle phase across phenotypic clusters in PDCLs.

**Supplementary Figure 20** Heatmap representations of therapy-induced plasticity by AMG232 and RT as recorded in PDCLs and high-grade glioma biopsies. (a) Percentage of responsive cells in each tumor-specific phenotypic cluster measured upon AMG232 treatment of the PDCL pool. (b) Percentage of responsive cells measured upon AMG232 treatment in each phenotypic cluster in the biopsy pool. (c) Percentage of responsive cells in each tumor phenotypic cluster measured upon RT treatment of the PDCL pool. (d) Percentage of responsive cells in each tumor phenotypic cluster measured upon RT treatment of patients' biopsies. (e) RT-driven enrichment/depletion of tumor phenotypic subclusters in PDCLs (normalized and z-scored). (f) RT-driven enrichment/depletion of tumor phenotypic subclusters in biopsies (normalized and z-scored). Samples in each heatmap are hierarchically clustered based on TP53 and (inferred) MDM2 mutational status.

#### Supplementary Tables:

**Supplementary Table 2** Antibody panels used for PDCLs and GBM samples.

| Target | Isotope | Clone | Vendor | PN | PDCLs | Biopsies |
|---|---|---|---|---|---|---|
| CD45 | 89Y | HI30 | Fluidigm | 3089003B | ○ | ● |
| GFAP | 112Cd | GA5 | ThermoFisher Scientific | 14-9892-82 | ● | ● |
| CC3 | 141Pr | C92-605 | BD Pharmingen | 559565 | ● | ● |
| Olig2 | 143Nd | 211F1.1 | Millipore | MABN50 | ● | ● |
| CD15 | 144Nd | W6D3 | Fluidigm | 3144019B | ○ | ● |
| CD31 | 145Nd | WM59 | Fluidigm | 3145004B | ○ | ● |
| MDM2 | 149Sm | 4H26L4 | ThermoFisher Scientific | 700555 | ● | ● |
| Sox2 | 150Nd | O30-678 | Fluidigm | 3150019B | ● | ● |
| Nestin | 151Eu | 25/Nestin | Fluidigm | 3151013A | ● | ● |
| CD44 | 153Eu | 691534 | Fluidigm | 3153021C | ● | ● |
| Vimentin | 154Sm | D21H3 | Fluidigm | 3154014A | ● | ● |
| p16 | 156Gd | 15C10C30 | BioLegend | 675602 | ● | ● |
| CD24 | 158Gd | REA832 | MACS Miltenyi Biotec | 130-124-316 | ● | ● |
| p21 | 159Tb | 12D1 | Fluidigm | 3159026A | ● | ● |
| PDGFRa | 160Gd | D13C6 | Fluidigm | 3159026A | ● | ● |
| Ki-67 | 161Dy | B56 | Fluidigm | 3161007B | ○ | ● |
| Ki-67 | 162Dy | B56 | Fluidigm | 3162012B | ● | ○ |
| CD11c | 162Dy | Bu15 | Fluidigm | 3162005B | ○ | ● |
| CCNB1 | 164Dy | GNS-1 | Fluidigm | 3164010A | ● | ● |
| p53 | 169Tm | DOI-1 | Abcam | ab1101 | ● | ● |
| CD3 | 170Er | UCHT1 | Fluidigm | 3170001B | ○ | ● |
| CD68 | 171Yb | KP1 | BioLegend | 916104 | ○ | ● |
| pEGFR | 146Nd | D7A5 | Fluidigm | 3146007A | ○ | ● |
| pH2A.X | 147Sm | JBW301 | Fluidigm | 3147016A | ● | ● |
| BAX | 165Ho | BAX/962 | Novus Biologicals Europe | NBP2-47815-0.1mg | ● | ● |
| pRB1 | 166Er | J112-906 | Fluidigm | 3166011A | ● | ● |
| pATM | 173Yb | 10H11.E12 | Abcam | ab36810 | ● | ● |
| pH3 | 175Lu | HTA28 | Fluidigm | 3175012A | ● | ● |
| CD11b | 209Bi | ICRF44 | Fluidigm | 3209003B | ○ | ● |

**Supplementary Table 3** Protein markers used to define phenotypic subpopulations of tumor cells

| Phenotypic ID | Markers |
|---|---|
| AC | GFAP+ |
| NPC | CD24+ NESTIN+ |
| MES | CD44+ VIMENTIN+ |
| OPC | OLIG2+ PDGFRa+ |
